# Antimicrobial nanolayers of thymol and carvacrol on titanium surfaces: the crucial role of interfacial properties in thymol’s superior osteogenic response

**DOI:** 10.1101/2024.08.30.610464

**Authors:** Ariel Gonzalez, Alejandro Miñán, Eduardo Prieto, Patricia Schilardi, Natalia S. Fagali, Mónica Fernández Lorenzo de Mele

## Abstract

“Green’’ nanotechnologies have emerged as environmentally friendly alternatives against microbial multidrug-resistant biofilms. In this study, bactericidal “green” nanolayers (NL) were developed on Ti surfaces using two isomeric phytocompounds, carvacrol (Carv-Ti-NL) and thymol (TOH-Ti-NL). These NLs were fabricated using a one-step immersion treatment method based on a simple and spontaneous self-assembly process. Both NLs revealed strong antimicrobial activity, displaying anti-biofilm and biocidal effects. Notably, TOH-Ti-NL exhibited superior osteogenic performance compared to Carv-Ti-NL, as evidenced by enhanced pre-osteoblast cell attachment and growth, and the production of ALP, collagen type I and Ca^2+^ deposition. In contrast, fibroblastic cells exhibited reduced attachment on TOH-Ti-NL and enhanced proliferation on Carv-Ti-NL. Considering the biological differential effects, the physicochemical properties of these conformational isomers’ NLs were studied to elucidate potential differences that could impact on cell response. Although the ATR-FTIR spectra of the NLs were similar and indicated the spontaneous oxidation of Carv and TOH leading to ketonic structures, distinct contributions were observed after the electrooxidation of each NL. Slight differences in hydrophilicity were found for both nanostructures, but higher roughness was found for TOH-Ti-NL. Furthermore, the release curves of Carv and TOH from the NLs revealed distinct profiles over time. Overall, Carv and TOH formed self-assembled layers on Ti able to eradicate *Staphylococcus aureus* biofilms. Their different physical and chemical characteristics induced distinct responses from eukaryotic cells attached to the NLs. Given these characteristics one might envisage the use of either Carv-Ti-NL or TOH-Ti-NL in order to fine-tune specific chemical physical properties of Ti-based implants.

**Graphical abstract:** 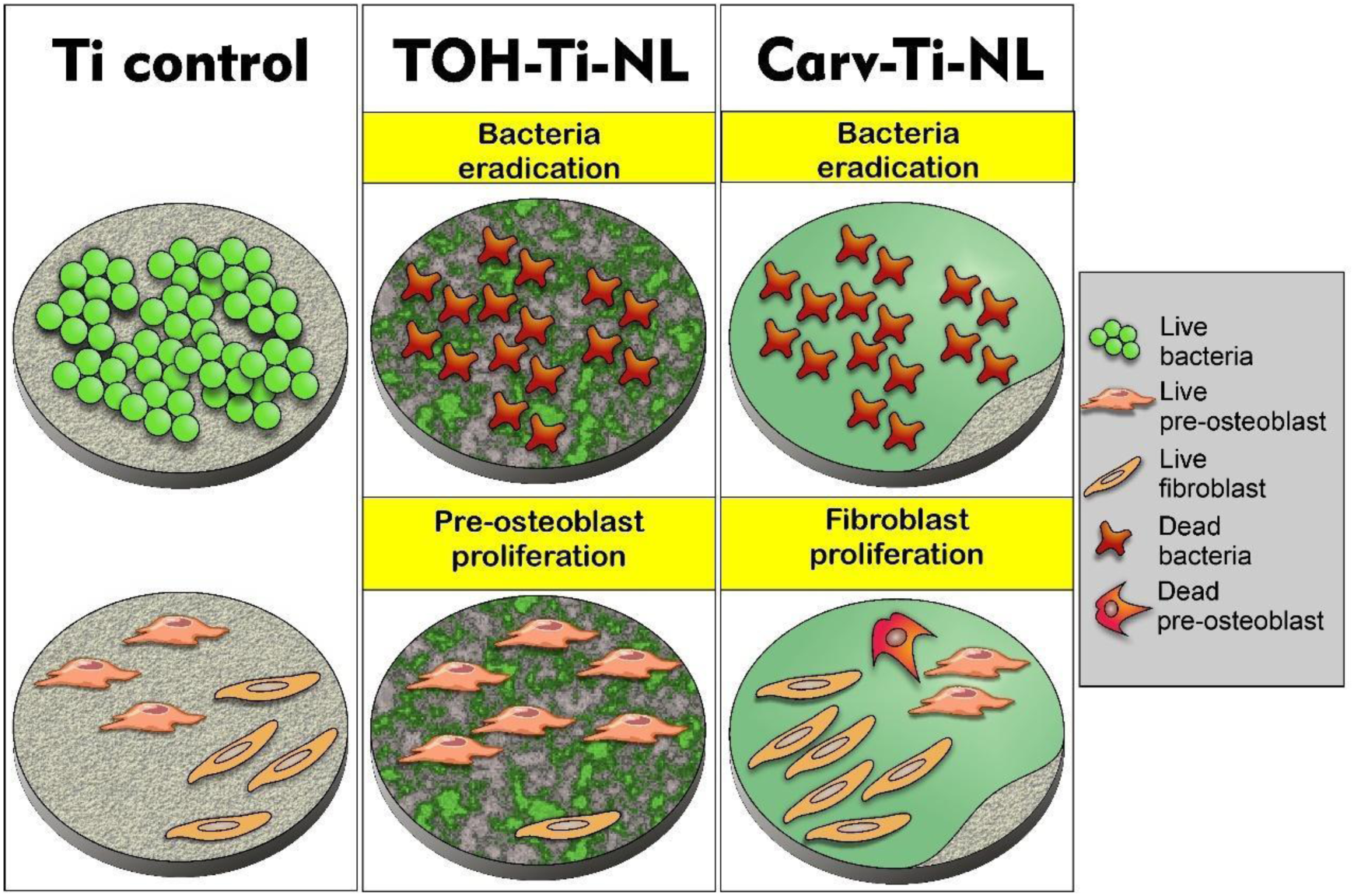

## 1. INTRODUCTION

Thymol (TOH) and carvacrol (Carv) are monoterpene phenols and conformational isomers, chemically known as 2-isopropyl-5-methylphenol and 5-isopropyl-2-methylphenol, respectively. Found abundantly in plant genera like *Thymus, Origanum,* and *Satureja*, these phenolic compounds (PhCs) have significant medical attributes. They are rapidly absorbed following oral administration and quickly degraded in the stomach or intestine [1]. PhCs exhibit diverse pharmacological properties, effective against cardiovascular, neurological, rheumatological, gastrointestinal, metabolic and malignant diseases. Their physicochemical properties, such as solubility, absorption or elimination rates, pose challenges in developing therapeutic strategies for systemic therapies [2]. Despite these barriers, various “green” nanomaterials have been developed to enhance drug delivery and other therapeutics application [3].

TOH and Carv are multifunctional compounds with notable therapeutic effects [4]. TOH is recognized for its anti-inflammatory, antioxidant and antihyperlipidemic properties. Both compounds aid in wound healing by modulating inflammatory cytokines and reducing oxidative stress. In the later stages, they promote re-epithelialization, angiogenesis, granulation tissue development, and collagen deposition, while also influencing fibroblast and keratinocyte growth [5]. PhCs are also well-known for their antimicrobial properties [6–10], showing efficacy against Gram-positive bacteria, such like *Staphylococcus aureus* [11,12], but are less effective against Gram-negative bacteria [13]. The antimicrobial action of Carv and TOH is attributed to their affinity for hydrophobic phases, allowing them to accumulate in bacterial membranes [14,15]. This accumulation causes membrane swelling, integrity loss, disruption of the proton motive force, and release of cellular components, leading to membrane destabilization [13,14,16]. Carv amphipathic nature also enables it to penetrate the polar polysaccharide matrix of biofilms, while TOH lipophilic characteristics allow it to interact with and damage phospholipid membranes [16–18]. Additionally, TOH can bind to genomic DNA of *S. aureus*, altering its structure and interfering with biological functions [19].

In a previous study [20], we demonstrated the effective antimicrobial action of TOH-coated implantable Ti. Infections associated with implantable biomaterials like orthopedic and dental implants pose significant challenges due to biofilm formation, which is resistant to conventional antibiotics [21–23]. Ti and its alloys are preferred for such implants due to their mechanical properties, corrosion resistance, and biocompatibility [24]. However, their susceptibility to microbial adhesion and biofilm formation has driven research into antimicrobial strategies for these materials. Beyond antimicrobial action, PhC-coatings must also exhibit osteogenic properties and ensure optimal interactions at the bone/implant interface. Natural compounds such as silibinin, resveratrol, quercetin, and genistein have demonstrated osteoinductive properties, promoting stem cell differentiation and extracellular matrix mineralization [25]. Essential oils containing Carv and TOH have been shown to enhance osteogenesis and adipogenesis of mesenchymal stromal cells *in vitro* [25].

This study aims to compare the biological responses of TOH and Carv on eukaryotic and prokaryotic cells using self-assembled nanolayers (NLs) formed on Ti (TOH-Ti-NL and Carv-Ti-NL, respectively) by a simple one-step immersion treatment method. We conducted a comparative analysis of their antimicrobial actions, biocompatibility and osteogenic properties, as well as the physicochemical and surface characteristics of the coatings. This analysis seeks to determine if there are distinct properties that influence specific biological responses, thereby refining the selection of suitable PhCs for self-assembled coatings, particularly those with similar molecular structures present in essential oils.

## 2. MATERIALS AND METHODS

### 2.1 Chemicals

TOH and Carv (Sigma, St. Louis, MO, USA) were used in the experimental procedures, and their respective structures are depicted in Fig. 1. Solutions were prepared using ultrapure water, while all chemical reagents employed in the trials were of analytical grade.

**Figure 1.**
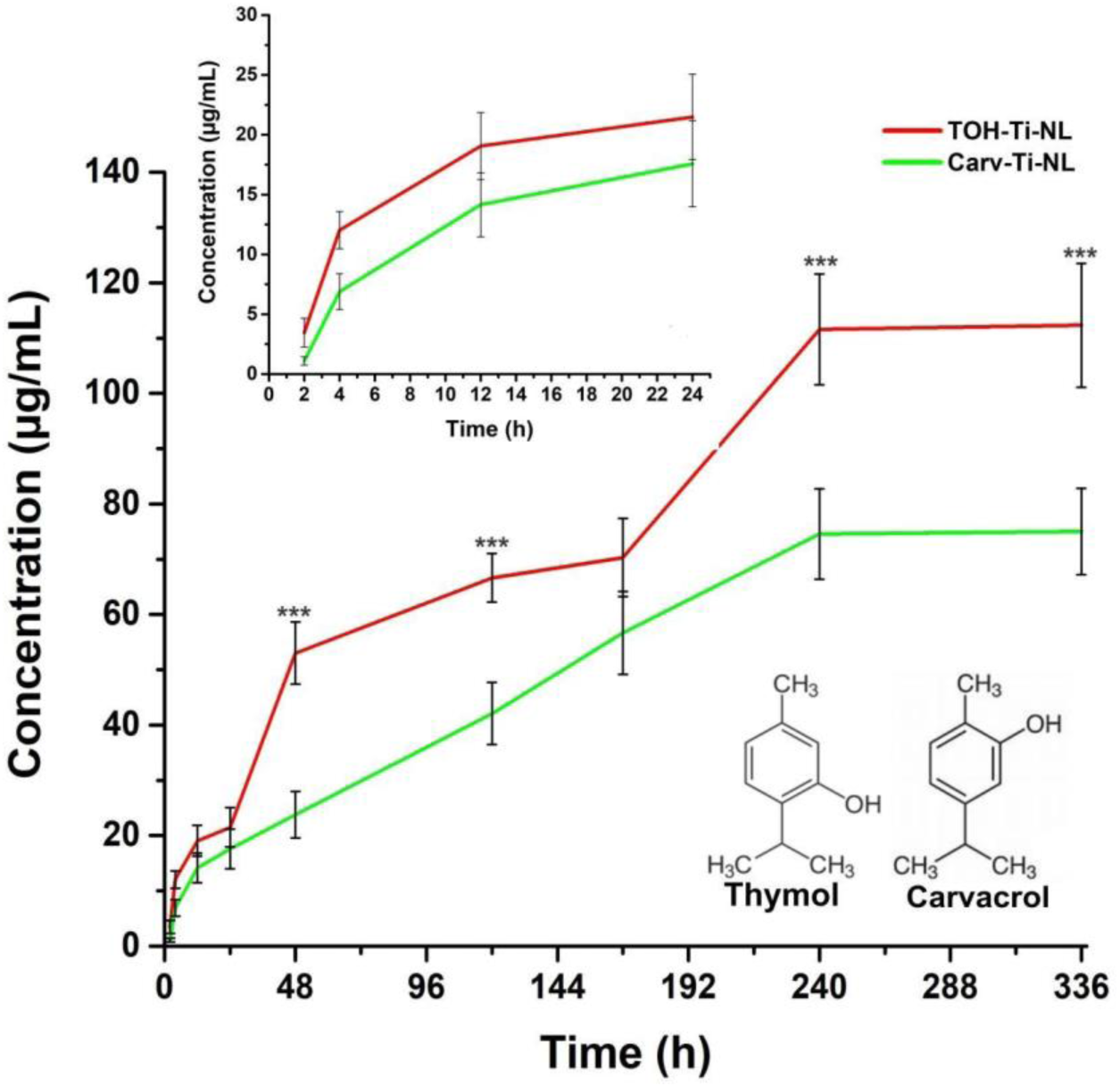
Release profiles from TOH-Ti-NL and Carv-Ti-NL. obtained by UV-Vis at λ = 274 and 273 nm, respectively, up to 14 days (Inset: release profile in the first 24 h). Molecular structures of TOH and Carv are shown. *** Indicates significant differences between TOH-Ti-NL and Carv-Ti-NL concentrations (p<0.001).

### 2.2 Titanium samples preparation

Cylindrical Ti disks (NMM Machinery Manufacturing Co. Ltd., Guangdong, China, 10 mm in diameter and 1 mm thick) were employed as substrate materials. The samples underwent a sequential process of mechanical polishing using P320, P400 and P600 grade SiC abrasive paper, followed by a subsequent chemical polishing step with a HF:HNO_3_:H_2_O solution (1:3:10) for 1 min. Afterwards, the Ti disks were thoroughly rinsed using ultrapure water and then used as0 supporting base for the deposition of TOH or Carv.

### 2.3 Formation of TOH or Carv nanolayers on Ti samples (TOH-Ti-NL and Carv-Ti-NL)

The formation of TOH-Ti-NL and Carv-Ti-NL involved a simple adsorption process onto the Ti surface. Initially, the polished Ti samples were immersed in 0.1M TOH or Carv ethanol/sulfuric acid solutions (30:70) for 2 hours, employing the one-step immersion treatment method. Then, the samples underwent thorough rinsing with Phosphate Buffered Saline Solution (PBS, pH 7.4) and ultrapure water for 1 min, aiming to eliminate weakly adsorbed TOH or Carv molecules. The resulting TOH-Ti-NL and Carv-Ti-NL samples were used in the subsequent assays. For comparative purposes, uncoated polished Ti disks were employed as controls (referred to as Ti control henceforth).

### 2.4 Release curves through UV-Vis spectroscopy

The release of adsorbed TOH or Carv molecules from TOH-Ti-NL or Carv-Ti-NL into the surrounding medium (PBS) was quantified using UV-Vis spectroscopy with a Shimadzu UV-1800 spectrophotometer at 274 nm to TOH and 273 nm to Carv [26]. The amounts of released TOH and Carv were measured at various time intervals up to 14 days (2, 4, 12, 24, 48, 120, 168, 240 and 336 h). Each assay was separately conducted in triplicate.

### 2.5 Microbiological tests

#### 2.5.1 Bacterial cultures

*Staphylococcus aureus* (ATCC 25923) was cultured by inoculating 100 mL of nutrient broth and then incubated overnight at 37 °C with agitation at 250 rpm. Following incubation, the number of microorganisms was adjusted to ∼10^8^ colony-forming units (CFU)/mL with fresh nutrient broth. To ensure the reliability of results, all antimicrobial assays detailed below were conducted a minimum of three times, confirming the reproducibility of the results.

#### 2.5.2 Inhibition halo assay of NLs

The antimicrobial activity against *S. aureus* of TOH-Ti-NL and Carv-Ti-NL was assessed using the agar diffusion test [27]. To this end, each sterile metal disk sample (TOH-Ti-NL, Carv-Ti-NL and Ti) were placed in the center of separated Petri dishes, seeded with *S. aureus* (100µL ∼ 10^8^ CFU/mL) and incubated at 37°C for 24 h. The diameters of the inhibition halos surrounding the metal samples were measured after the incubation period. Ti samples were used as negative controls to check the absence of inhibition halo formation.

#### 2.5.3 Inhibitory effect of NLs on bacterial adhesion

The inhibitory effect against initial bacterial adhesion on Ti control and on recently formed TOH-Ti-NL and Carv-Ti-NL layers was assessed. With this purpose, each sterile sample was placed within a well of a 6-well culture plate. Subsequently, 8 mL of a *S. aureus* suspension (∼10^8^ CFU/mL) was poured on each substrate followed by incubation for 3h at 37 °C. The weakly adhered bacteria were removed by washing. Then, bacteria firmly attached to the surfaces were removed through sonication for 10 min. The number of detached bacteria was quantified using the serial dilution method and CFUs plate counting [27].

#### 2.5.4 Antimicrobial effects of NLs against biofilms

##### 2.5.4.1 Antimicrobial effect of recently formed NLs against biofilms

The antimicrobial activity against biofilms (24 h biofilm formation) that grew on recently formed TOH-Ti-NL, and Carv-Ti-NL and on Ti control surfaces was investigated by immersing the samples in a *S. aureus* suspension (∼10^8^ CFU/mL). After the 24 h incubation period, bacteria that were firmly adhered were removed by sonication, and the count of viable bacteria was determined through serial dilution followed by CFUs counting [20].

##### 2.5.4.2 Remaining antimicrobial activity of NLs after 24 h and 48 h release periods against biofilms formation

Tests were conducted to determine the degree of antimicrobial activity present after the TOH-Ti-NL and Carv-Ti-NL samples were immersed in sterile nutrient broth for 24 and 48 h. To assess the remaining antimicrobial activity, the samples were immersed in this medium to allow the release of organic compounds from the NLs. Subsequently, they were incubated with a bacteria suspension following the procedure described in 2.5.4.1 (24 h incubation for biofilm development).

#### 2.5.5 Live/Dead staining

Live/Dead BacLight® (Invitrogen) staining was employed to assess the viability of bacteria attached to TOH-Ti-NL, Carv-Ti-NL and Ti control surfaces after the completion of experiments described in sections 2.5.3 and 2.5.4. The staining was carried out by following the manufacturer protocol. After staining, the surfaces were imaged by epifluorescent microscopy. The outcomes were presented as the percentage of the area covered by live/dead bacteria (%) on TOH-Ti-NL and Carv-Ti-NL surfaces, relative to the results obtained from the Ti control surface according to [27]:

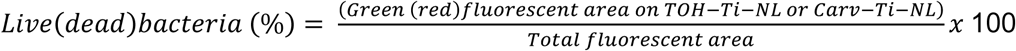

#### 2.5.6 Evaluation of S. aureus extracellular polymeric matrix formation on TOH-Ti-NL and Carv-Ti-NL substrates

To assess the formation of the extracellular polymeric matrix (EPM) generated by biofilms on TOH-Ti-NL, Carv-Ti-NL and Ti control surfaces, Sypro Ruby^(R)^ (Invitrogen) staining was utilized after the completion of the assays described in section 2.5.4. The extent of EPM production on TOH-Ti-NL and Carv-Ti-NL surfaces was quantified as the percentage of fluorescent covered area attributed to the polymeric matrix, relative to the Ti control according to [28]:

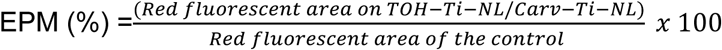

### 2.6 Cytocompatibility and osteogenic activity of TOH-Ti-NL and Carv-Ti-NL

#### 2.6.1 Cell cultures

Two cell lines from *Mus musculus* were employed: fibroblasts (L929, ATCC, USA) and pre-osteoblasts (MC3T3-E1, ATCC, USA). The cultures were maintained at 37°C in a humid atmosphere containing 5% CO_2_.

#### 2.6.2 Cytotoxic effect of NLs extracts by reduction of methyl tetrazolium (MTT)

The effect of NLs extracts on metabolic activity was assessed through the MTT assay on L929 and MC3T3-E1 cell lines according to ISO 10993-5:2017 and previously established protocols [20]. Initially, extracts were obtained following a 24 h-immersion of TOH-Ti-NL, Carv-Ti-NL and Ti control in DMEM culture medium (without cells). Subsequently, 20.000 cells/well were seeded in a 96-multiwell dish, incubated for 24 h, and then treated for 24 h with the extracts. Following this, the media were replaced, and the cells were incubated with 0.5 mg/mL MTT (3-(4,5-dimethylthiazol-2-yl)-2,5-diphenyl-tetrazolium-bromide, Sigma–Aldrich, St. Louis, MO, USA) under standard culture conditions for 3 h. Cell viability was monitored by the conversion of MTT to the colored formazan product through mitochondrial dehydrogenase activity. The colored product was measured in a Microplate Reader (QuantBioTek, USA) at λ= 570 nm after cell lysis in DMSO (100 μL/well). Mitochondrial activity was expressed as a percentage of the Ti control values.

#### 2.6.3 Cell adhesion and proliferation on NLs

Fibroblastic and pre-osteoblastic cells (30.000 cells/mL) were seeded on TOH-Ti-NL, Carv-Ti-NL and Ti control surfaces placed in 24-well plates, and were incubated for 1, 2, 5 and 7 days. Following the incubation period, cell adhesion and proliferation were evaluated using acridine orange staining. The percentage of attached cells on TOH-Ti-NL and Carv-Ti-NL surfaces was calculated by comparing the fluorescent area covered by cells in these samples to the area of cells attached on Ti control sample (1 day).

#### 2.6.4 Osteogenic activity of NLs

##### 2.6.4.1 Osteogenic cell differentiation

To evaluate the osteogenic activity of TOH-Ti-NL, Carv-Ti-NL and Ti control surfaces, pre-osteoblastic MC3T3-E1 cells were seeded at 30.000 cell/mL onto the surface samples. These cells were then incubated with an osteogenic medium following established protocol [29,30].

##### 2.6.4.2 Semi-quantitative determination of Alkaline Phosphatase Activity

After 15 days of incubation in osteogenic medium, cells attached on TOH-Ti-NL, Carv-Ti-NL and Ti control surfaces were rinsed with a PBS solution and then stained with Vector Red Alkaline Phosphatase (ALP) Substrate kit [31]. The ALP activity was semi-quantitatively calculated and expressed as the percentage of the fluorescent area of each surface observed by epifluorescent microscopy relative to the Ti control.

##### 2.6.4.3 Quantitative determination of type I collagen production

After 21 days of incubation in the osteogenic medium, the production of type I collagen was determined using the Sirius Red colorimetric assay [31]. Briefly, cells attached on Ti samples were washed with PBS and fixed with Bouin solution. Next, the samples were colored with Sirius Red solution for 1 h. Finally, the colored product was measured in a Microplate Reader (QuantBioTek, USA) at λ= 550 nm after cell lysis with NaOH 1N. Collagen production for each sample was expressed as a percentage relative to the Ti control.

##### 2.6.4.4 Quantitative determination of mineralization

After 21 days of incubation with osteogenic medium, the mineralization degree was determined by quantifying the deposition of Ca (II) using the Alizarin Red colorimetric assay [31]. Shortly, the cells attached on Ti samples were washed with PBS and fixed with 10% of formol. Then, the samples were colored with alizarin red for 10 min. Finally, the colored product was measured in a Microplate Reader (QuantBioTek, USA) at λ= 548 nm after cell lysis with NaOH 1N. The degree of mineralization for each sample was expressed as a percentage relative to the Ti control.

##### 2.6.4.5 Quantification of proteins

The quantification of proteins as an indicator of cell population on TOH-Ti-NL, Carv-Ti-NL and Ti control surfaces was performed using the Bradford method [32] after differentiation in the osteogenic medium. Briefly, cells attached on Ti samples were washed with PBS and lysed with 0.1% Triton X-100. Next, the cell suspensions were centrifuged at 12.000 rpm for 15 min at 4°C. Afterwards, the supernatant with protein suspensions were collected and then, aliquots of them were placed in 96-well plates with Bradford reagent for protein determination. Absorbance was measured at λ = 595 nm, and standard curve was performed using bovine serum albumin (BSA) in the 0 to 20 µg/mL range. The protein quantity of the samples was expressed as a percentage relative to the Ti control.

### 2.7 Physicochemical characterization of TOH-Ti-NL and Carv-Ti-NL

#### 2.7.1 Attenuated Total Reflectance Fourier-Transform Infrared spectroscopy (ATR-FTIR)

ATR-FTIR spectra of the pure TOH and Carv and of TOH-Ti-NL and Carv-Ti-NL samples recently prepared were obtained. Additionally, spectra were taken after the anodic polarization (AP) of TOH-Ti-NL and Carv-Ti-NL in the (−1.0V, +2.0V) potential range at 50 mV/s (see 2.7.3.2 section). Spectra were obtained using a Varian 660 spectrometer equipped with an attenuated total reflection (ATR) accessory (MIRacle ATR, Pike technologies) featuring a ZnSe prism. In all instances, each ATR-FTIR spectrum resulted from the accumulation of 256 scans taken at a resolution of 2 cm^−1^.

#### 2.7.2 Atomic Force Microscopy (AFM) imaging

AFM was employed to perform topographic analysis of the samples. Images were acquired in tapping mode using a Nanoscope V microscope (Bruker, Santa Barbara, CA) with silicon tips (Arrow® NCR; NanoWorld, Neuchâtel, Switzerland). The imaging was carried out over a 2 x 2 μm^2^ area, and the surface roughness parameters were calculated using NanoScope Analysis 2.0 software [27]. The calculated parameters are defined as follows: Rq, root mean square average of height deviations, taken from the mean image data plane; Ra, arithmetic average of the absolute values of the surface height deviations measured from the mean plane; Rmax, maximum vertical distance between the highest and lowest data points in the image; Ssk (skewness, asymmetric parameter) measures the symmetry of surface data about a mean data profile; Sku (kurtosis parameter) indicates whether data are arranged flatly or sharply about the mean. Rsa% (% of image surface), represents the relationship between the total area and the projected area in the image, reporting the changes that the nanofilms produce in the nanoroughness.

#### 2.7.3 Electrochemical measurements

##### 2.7.3.1 Open Circuit Potential measurements

To determine the electrochemical behavior of TOH and Carv during the formation of NLs, the open circuit potential (OCP) was measured for a 2h period [33]. This measurement was performed using ethanol/sulfuric acid solutions (30:70) without and with 0.1M TOH or Carv as the electrolyte. Ti samples were used as working electrodes, Pt electrode was employed as a counter electrode and saturated calomel electrode (SCE) was used as reference.

##### 2.7.3.2 Cyclic voltammetry and Anodic Polarization

The stability of the coatings was assessed through cyclic voltammetry (CV) in a 5 mM KCl electrolyte solution. The measurement involved two successive cycles at a 50 mV/s scan rate between −1.0 V and +2.0 V vs SCE reference electrode. Prior to each electrochemical run, the OCP was measured 1 min after immersion [34]. Occasionally, an anodic polarization (AP) in the −1.0 V to 2.0 V vs SCE potential range was made to force the electrooxidation process and evaluate the composition of the NLs through ATR-FTIR after this treatment.

##### 2.7.3.3 Tafel curves

The corrosion potential of TOH-Ti-NL or Carv-Ti-NL was evaluated through Tafel curves. Potentiodynamic curves were recorded at 1 mV/s scan rate, within the potential range from the OCP to −1 V in the cathodic direction and from the OCP to 1.0 V in the anodic direction. A 5 mM KCl was used as the electrolyte solution.

### 2.8 Statistical analysis

All results were expressed as mean ± standard error, and statistical differences were analyzed using ANOVA followed by the Multiple Range Test of Bonferroni with the GraphPad Prism 5.0 software.

## 3 RESULTS

### 3.1 Antimicrobial activity of TOH-Ti-NL and Carv-Ti-NL

The antimicrobial effect of TOH-Ti-NL and Carv-Ti-NL (see molecular structures in Fig. 1) substrates against *S. aureus* was evaluated through a series of experiments with focus on the inhibition of bacterial growth and surface adhesion, the formation of biofilms and the remaining antimicrobial activity of the substrates over time.

#### 3.1.1 Bacterial growth inhibition and release profiles of TOH and Carv from NLs

The diameters of the inhibition halos were measured after 24 h of incubation (Table 1) and the results indicated that TOH-Ti-NL exhibited a slightly larger bacterial inhibition halo than Carv-Ti-NL (2.0±0.2 mm and 1.8±0.2 mm, respectively). However, no statistically significant differences were found. As expected, the Ti control did not display any inhibition halo.

**Table 1.**
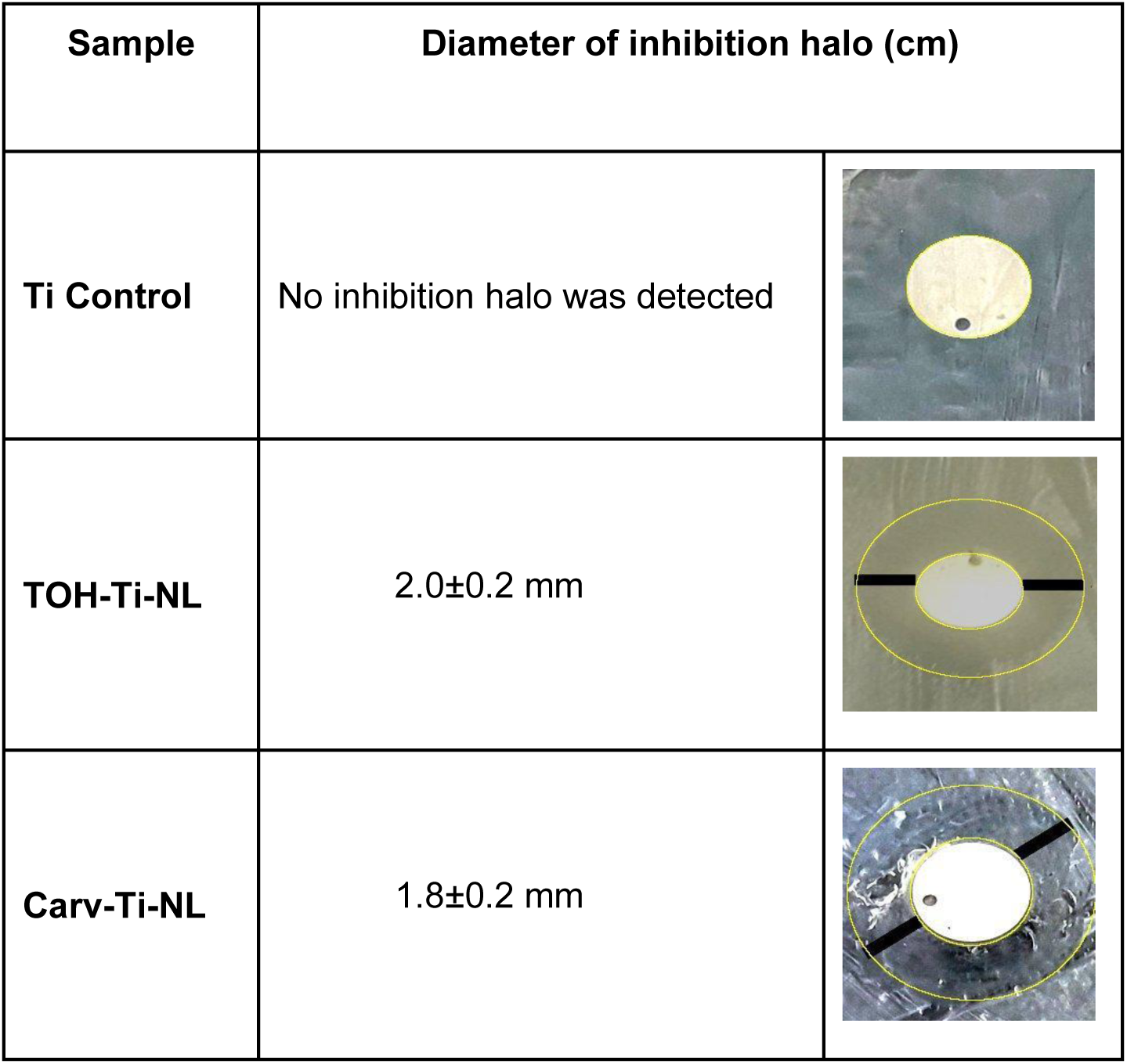
Measurements of inhibition halo.

| Sample | Diameter of inhibition halo (cm) |  |
| --- | --- | --- |
| **Ti Control** | No inhibition halo was detected |  |
| **TOH-Ti-NL** | $2.0\pm0.2$ mm | |
| **Carv-Ti-NL** | $1.8\pm0.2$ mm | |

To interpret these results, it is useful to investigate the release profiles of both NLs (Fig. 1). Pure TOH and Carv display similar UV-Vis spectra, with the maximum absorbance occurring at 274 and 273 nm respectively (Fig. 1S). The release curves from TOH-Ti-NL and Carv-Ti-NL in PBS solution were quantitatively determined through absorbance at those wavelengths. The release profile, within the first 24 h, indicated that the average cumulative concentration of products measured from Carv-Ti-NL samples were slightly lower than those of TOH-Ti-NL (Fig. 1, inset), although the differences were not statistically significant. During this period, both NLs exhibited a high initial release rate like a burst release (average rate: 0.90±0.04 μg/h and 0.73±0.02 μg/h for TOH-Ti-NL and Carv-Ti-NL, respectively, which decreased as 24 h was reached. Inhibition halo results (Table 1) are consistent with those obtained for the quantification of the release of products from NLs, where the release profile of TOH and Carv did not exhibit significant differences during the first 24 h.

Subsequently, the release of Carv occurred at a constant rate (average rate: 0.31±0.02 μg/h) for up to 10 days (240 h), with the Carv concentration remaining unchanged thereafter. In contrast, the TOH release curve displayed three phases: the already mentioned up to 24 h (Fig. 1, inset) (average rate: 0.90±0.04 μg/h), a second increase in TOH concentration between 24 and 48 h (average rate: 1.3±0.1 μg/h) followed by a plateau up to 168 h (average rate: 0.14±0.02 μg/h) and, finally, a third increase until the 10th day (240 h). No further release was observed from 240 h to 336 h (14 days).

#### 3.1.2 TOH-Ti-NL and Carv-Ti-NL inhibitory effect on bacterial adhesion (3h of incubation)

The bacterial adhesion was assessed on the Ti control, TOH-Ti-NL and Carv-Ti-NL surfaces following a 3h incubation period with *S. aureus*. The results of the CFUs counting are presented in Fig. 2A. Bacterial adhesion was observed on the Ti control substrate, reaching (4.0±0.5) x10^5^ CFU/cm^2^. In contrast, no viable bacteria were found on the TOH-Ti-NL and Carv-Ti-NL substrates. Furthermore, fluorescence microscopy using the Live/Dead staining technique demonstrated that, after 3h, bacteria were attached to the TOH-Ti-NL and Carv-Ti-NL surfaces, but they were completely dead, while those on the Ti control remained viable (Fig. 3A, B).

**Figure 2.**
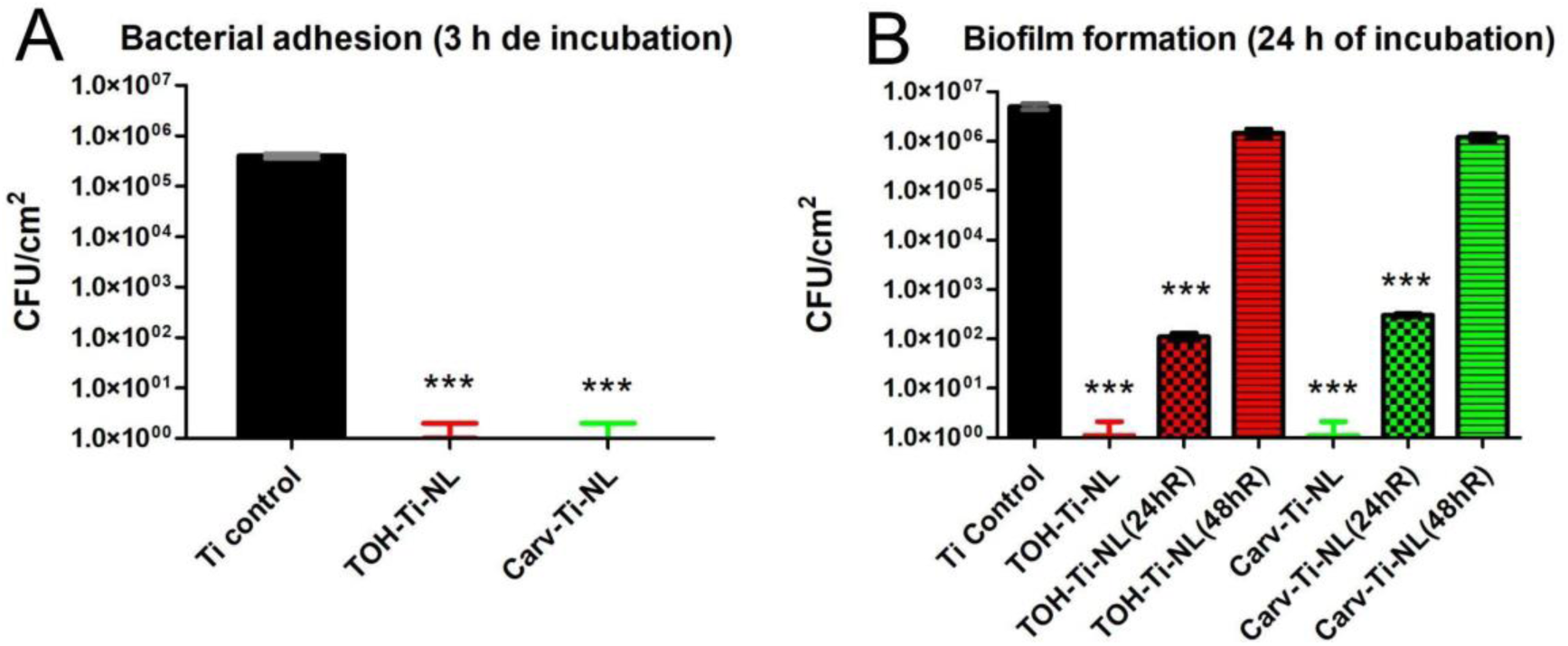
Antimicrobial activity of NLs against *S. aureus:* Serial dilution and CFU counting method. **(A)** Inhibitory effect of TOH-Ti-NL (red) and Carv-Ti-NL (green) on bacterial adhesion (3h in bacterial culture). **(B)** Effect of NLs against biofilms (24 h in bacterial culture) developed on: i) recently formed TOH-Ti-NL and Carv-Ti-NL; ii) the remaining NLs after 24 h-release period (TOH-Ti-NL(24hR), Carv-Ti-NL(24hR)) and iii) after 48 h-release period (TOH-Ti-NL(48hR), Carv-Ti-NL(48hR)). A polished Ti disk was used as a control. *** Indicates significant differences with Ti control samples (p<0.001).

**Figure 3.**
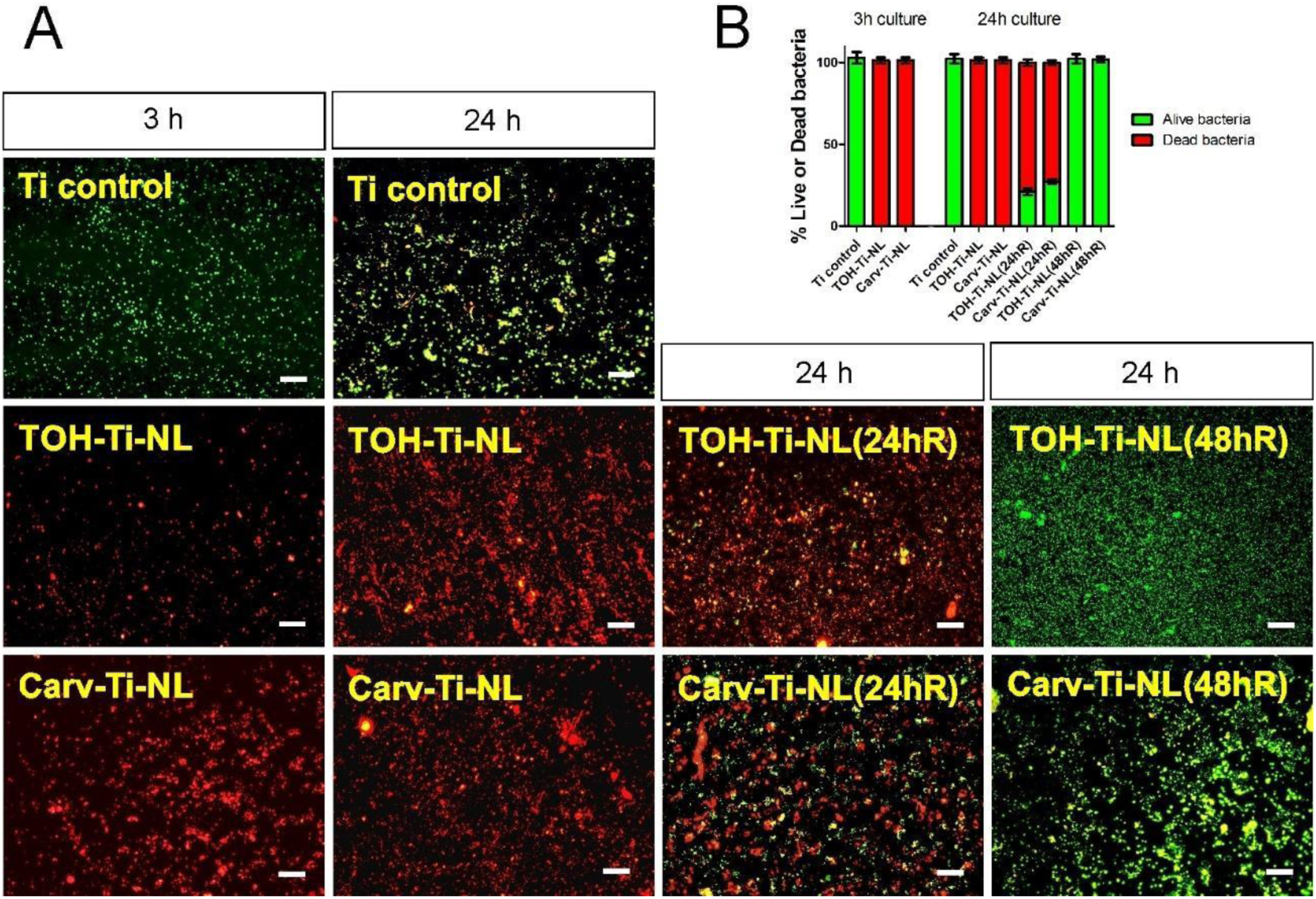
Antimicrobial activity of NLs against *S. aureus*: Live/Dead staining. **(A)** Live/Dead staining images (Green: live cells, Red: dead cells, 40X objective, scale bar: 20 µm) and **(B)** quantification of Live/dead staining of bacteria adhered (3 h in bacterial culture) and biofilm (24 h in bacterial culture) developed on i) recently formed TOH-Ti-NL and Carv-Ti-NL, ii) the remaining NLs after 24 h -release period (TOH-Ti-NL(24hR) or Carv-Ti-NL(24hR) and iii) 48 h-release period (TOH-Ti-NL(48hR) or Carv-Ti-NL(48hR). All results are expressed as a percentage of Ti control samples. A polished Ti disk was used as a control.

#### 3.1.3 Antimicrobial effect of recently formed Carv-Ti-NL and TOH-Ti-NL against biofilms

To investigate the effect of TOH-Ti-NL and Carv-Ti-NL on biofilms, these samples along with a Ti-control were exposed to a *S. aureus suspension* for 24 h. The biofilm formation was evaluated using Live/Dead staining and enumerated by the plate counting method. Additionally, the production of EPM generated by early biofilms was determined using Sypro-Ruby staining.

After 24 h, the Ti control showed (5.0±0.7) x10^6^ CFU/cm^2^, while no viable bacteria were observed on TOH-Ti-NL and Carv-Ti-NL (Fig. 2B). Live/Dead staining revealed that the colonies grown on the Ti control were alive, whereas those attached to TOH-Ti-NL and Carv-Ti-NL were mostly dead (Fig. 3A, B). Furthermore, Sypro Ruby staining showed that the production of EPM significantly decreased in the presence of TOH or Carv NLs (>95% reduction compared to the Ti control) (Fig. 4).

**Figure 4.**
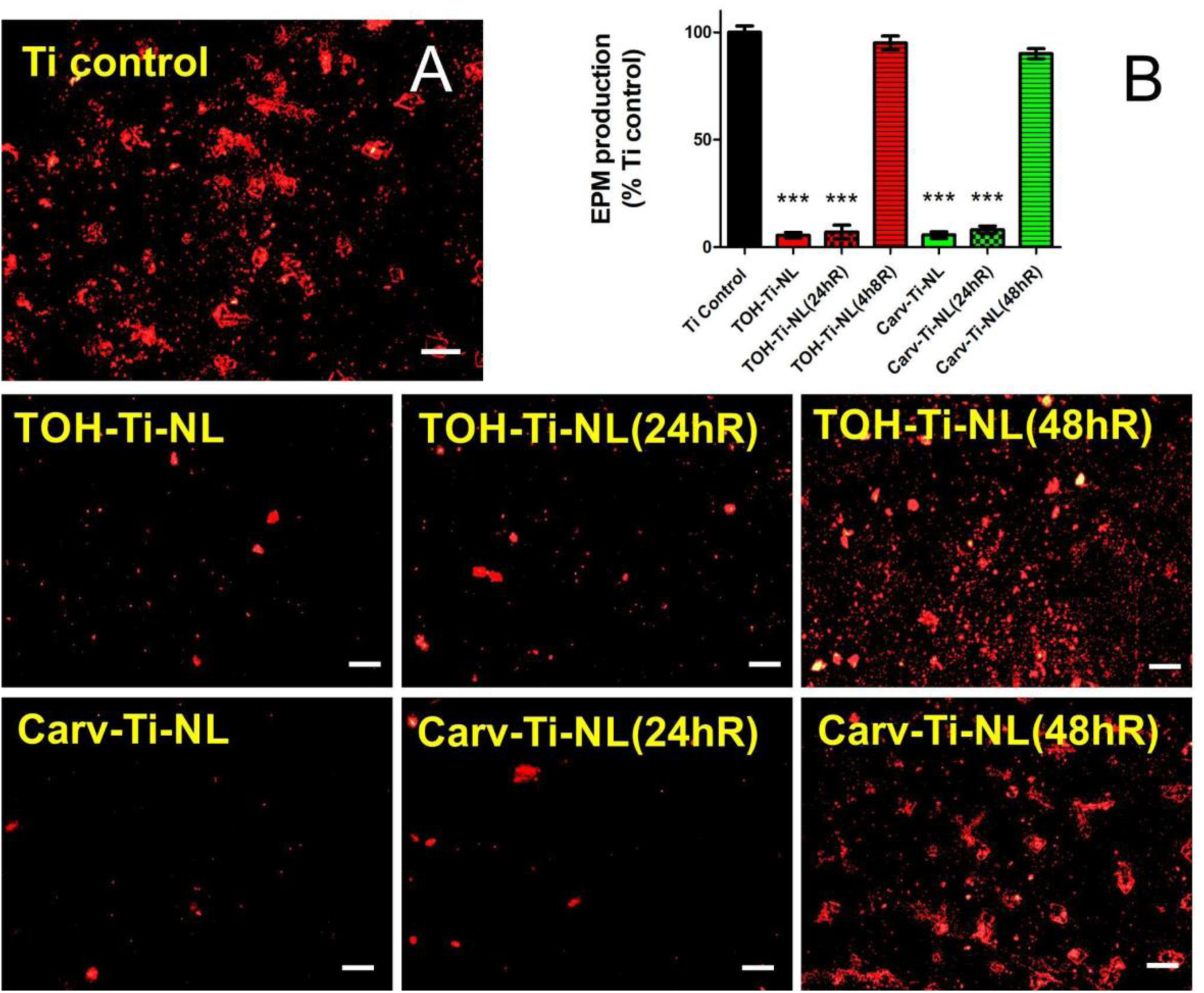
Antimicrobial activity of NLs against *S. aureus:* EPM production. (the higher red staining area the greater the EPS production). **(A)** Epifluorescence images of Sypro Ruby staining (10X objective, scale bar: 100 µm) and **(B)** quantification of EPM production of biofilm (24 h in bacterial culture) developed on i) recently formed TOH-Ti-NL and Carv-Ti-NL; ii) the remaining NLs after 24 h-release period (TOH-Ti-NL(24hR) or Carv-Ti-NL(24hR)) and iii) 48 h-release period (TOH-Ti-NL(48hR) or Carv-Ti-NL(48hR)). *** Indicates statistically significant differences compared to polished Ti control (p<0.001).

#### 3.1.4 Remaining antimicrobial effect of TOH-Ti-NL and Carv-Ti-NL after 24 h and 48 h release periods

To assess the remaining antimicrobial effect of TOH-Ti-NL and Carv-Ti-NL following different release periods, samples were initially exposed to sterile nutrient broth for 24 and 48 h. Subsequently, the 24 h-remaining-NLs-samples (*TOH-Ti-NL(24 hR)* and *Carv-Ti-NL(24hR)*) and 48-remaining-NLs-samples (*TOH-Ti-NL(48hR)* and *Carv-Ti-NL(48hR)*) were exposed to a *S. aureus* culture for an additional 24 h-period. The bacterial viability and the EPM production by the biofilms on these samples were evaluated by Live/Dead staining and CFU quantification respectively.

After a 24 h-release period, (1.1±0.2) x10^2^ and (3.0±0.3) x10^2^ CFUs were counted on TOH-Ti-NL(24hR) and Carv-Ti-NL(24hR) samples, respectively (Fig. 2B). This indicates that the bacterial viability on the samples decreased by four orders of magnitude compared to the Ti control (5.0±0.7) x10^6^, demonstrating a bactericidal effect of NLs even after 24 h of release. Live/Dead staining (Fig. 3) revealed that most of the attached bacteria on the TOH-Ti-NL(24hR) and Carv-Ti-NL(24hR) samples were dead. Sypro Ruby staining (Fig. 4) revealed a significant decrease in the production of the EPM on TOH-Ti-NL(24hR) and Carv-Ti-NL(24hR) samples compared to Ti control.

After a 48 h-release period, TOH-Ti-NL(48hR) and Carv-Ti-NL(48hR) samples showed bacteria viability (Fig. 2B), EPM production (Fig. 4) and percentage of live bacteria (Fig. 3) similar to those of the Ti control. These results indicate that after a 48 h-release period, the NLs lost their antimicrobial activity against *S. aureus,* although a significant release of TOH and Carv is still observed from NLs (Fig. 1).

### 3.2 Effect of TOH-Ti-NL and Carv-Ti-NL on eukaryotic cells

Fibroblast and pre-osteoblast cell attachment and proliferation were assessed to evaluate the cytocompatibility of TOH-Ti-NL and Carv-Ti-NL, through acridine orange staining in 1 to 7-day assays. The areas covered by cells were measured and the results were expressed as % of control Ti (Fig. 5).

**Figure 5.**
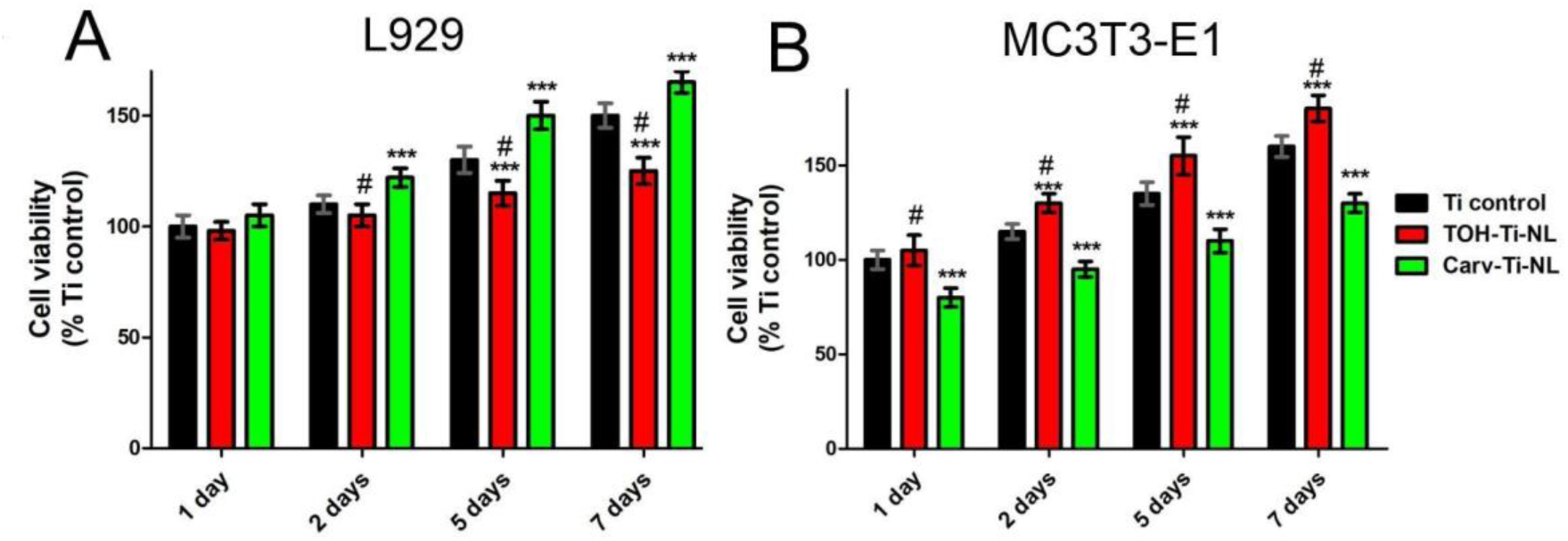
Cytocompatibility of NLs determined by acridine orange staining. **(A)** Fibroblastic cell (L929) adhesion and proliferation on NLs up to 7 days. **(B)** Pre-osteoblastic cells (MC3T3-E1) adhesion and proliferation on NLs up to 7 days. Cells were stained with Acridine Orange. Covered areas were measured and the results were expressed as % of Ti control 1 day. *** Indicates statistically significant differences compared to polished Ti control at each exposure period and # indicates significant differences between TOH-Ti-NL and Carv-Ti-NL (p<0.001).

The area covered by fibroblasts on TOH-Ti-NL samples was similar to that on Ti control at 1 and 2 days. However, at 5 and 7 days TOH-Ti-NL samples decreased fibroblastic cells proliferation (Fig. 5A). In contrast, pre-osteoblast cells exhibited covered areas larger than Ti control from 2 to 7 days, in consequence, TOH-Ti-NL increased pre-osteoblastic cell proliferation (Fig. 5B). Conversely, from 2 days, the area covered by fibroblastic cells was greater on Carv-Ti-NL than on the Ti control. However, pre-osteoblastic cells showed less affinity for Carv-Ti-NL with approximately 20-25% less area covered compared to the Ti control for all evaluated time points.

The cytotoxic effects of 24 h-NLs-extracts on pre-osteoblastic and fibroblastic cells were assessed using the MTT assay. The findings revealed that the TOH-Ti-NL extract had a non-cytotoxic effect on both fibroblastic and pre-osteoblastic cells. Conversely, the Carv-Ti-NL extract demonstrated a significant cytotoxic effect on pre-osteoblastic cells but none on fibroblastic cells (Fig. 6).

**Figure 6.**
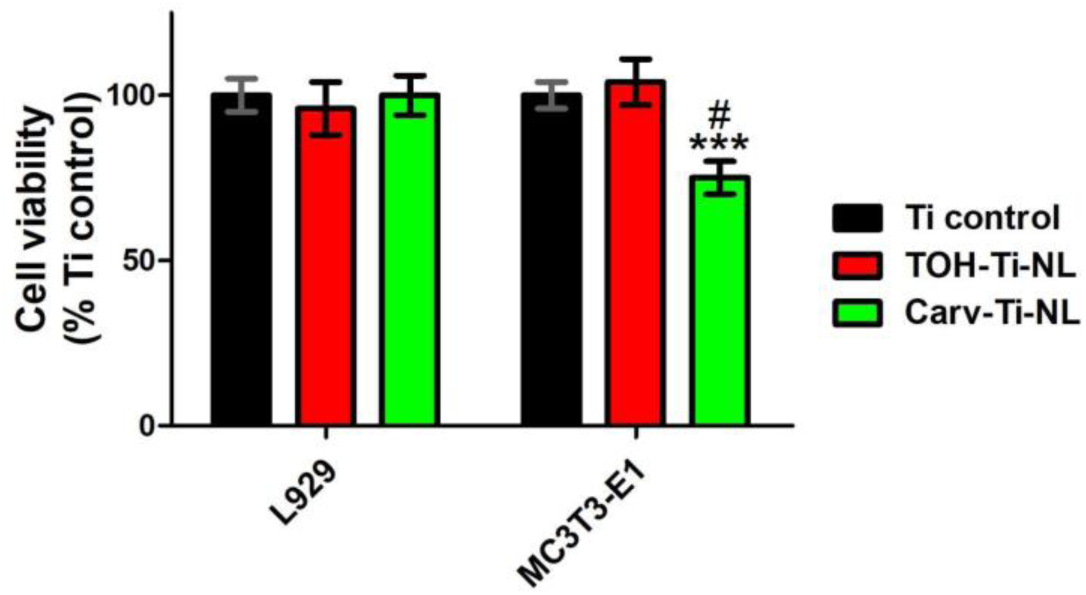
Cytotoxicity of NLs evaluated on fibroblastic (L929) and pre-osteoblastic (MC3T3-E1) cells by MTT assay. (***) Indicates statistically significant differences compared to polished Ti control and (#) indicates significant differences between TOH-Ti-NL and Carv-Ti-NL (p<0.001).

To evaluate the osteogenic differentiation activity of TOH- and Carv-NLs, the production of ALP, type I collagen, Ca^2+^ deposition and total protein content were assessed in differentiated pre-osteoblastic cells cultured onto these surfaces (Fig. 7).

**Figure 7.**
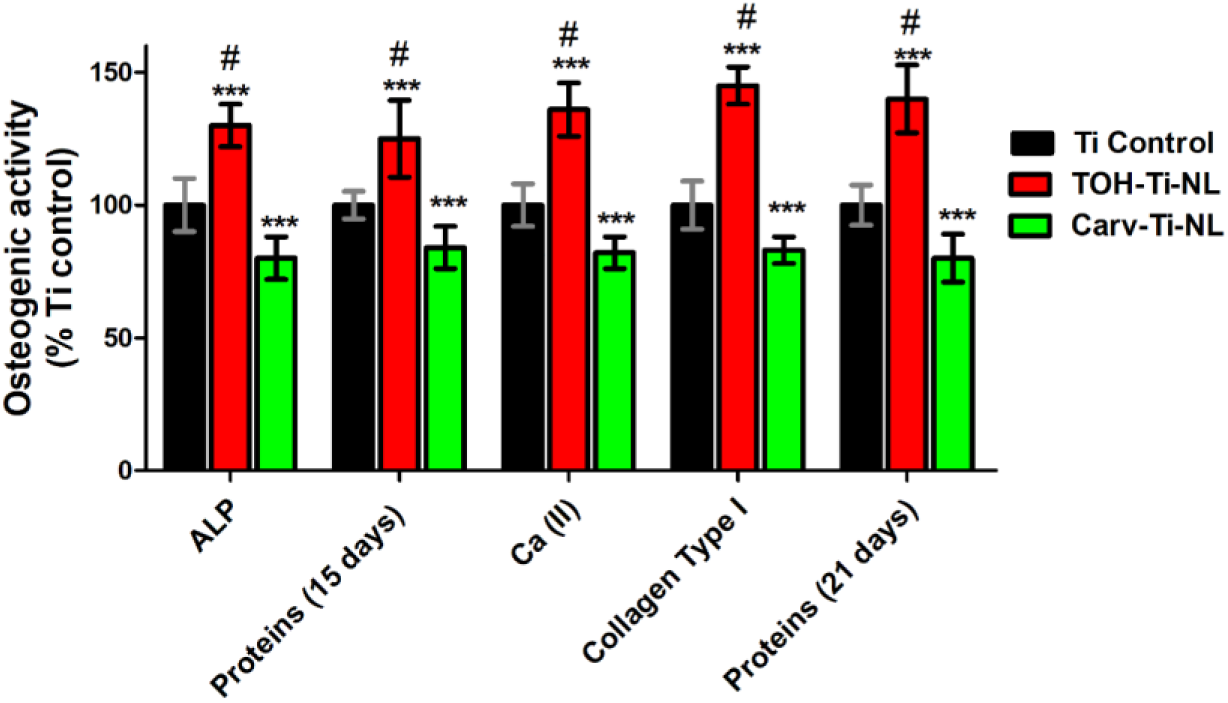
Osteogenic activity of NLs assessed at 15 and 21 days of incubation with pre-osteoblastic cells in the osteogenic medium. (***) Indicates statistically significant differences compared to Ti control and (#) indicates significant differences between TOH-Ti-NL and Carv-Ti-NL (p<0.001).

On TOH-Ti-NL samples, the production of ALP, Ca^2+^ and collagen type I increased by approximately 30%, 35% and 45%, respectively, compared to the Ti control. Additionally, total protein content showed an approximately 25 and 30% increase, at 15 and 21 days respectively, compared to Ti control samples. These results indicate that the population of differentiated cells, and, in consequence, the osteogenic parameters, increased on TOH-Ti-NL compared to Ti control. In contrast, results corresponding to Carv-Ti-NL demonstrated a decrease of approximately 20% in all these parameters compared to the Ti control. This suggests a lower population of differentiated cells on this surface, which could be explained due to Carv-Ti-NL samples producing a cytotoxic effect on pre-osteoblastic cells.

### 3.3 Physicochemical analysis

To assess the influence of the physicochemical properties of the NLs on the biological response, several analyses as ATR-FTIR, wettability, surface roughness by AFM images, and electrochemical assays of TOH-Ti-NL and Carv-Ti-NL were conducted.

#### 3.3.1 ATR-FTIR analysis

##### 3.3.1.1 ATR-FTIR spectra of pure compounds, TOH-Ti-NL and Carv-Ti-NL

ATR-FTIR spectra of pure TOH, pure Carv, TOH-Ti-NL and Carv-Ti-NL are shown in Fig. 8. The characteristic contributions of pure TOH and Carv were previously reported by several authors [35–38]. Pure TOH and Carv showed typical contributions from the aromatic ring due to the C-H stretching (3100-2850 cm^-1^), C=C stretching (1600-1500 cm^-1^) together with C-H out-of-plane bending (below 900 cm^-1^). The -OH stretching broad peak (3378 cm^-1^) in the liquid pure Carv corresponds to a -OH group involved in hydrogen bond with other Carv molecules, however pure solid TOH presents a narrower peak at 3171 cm^-1^ because this group is blocked in the solid crystal. The isopropyl group is detected in both compounds through a peak near 1420 cm^-1^. Pure Carv and pure TOH also present various peaks owed to C-O stretching (1300-1000 cm^-1^) with a specific peak owed to C-O stretching in phenols (1200-1260 cm^-1^).

**Figure 8.**
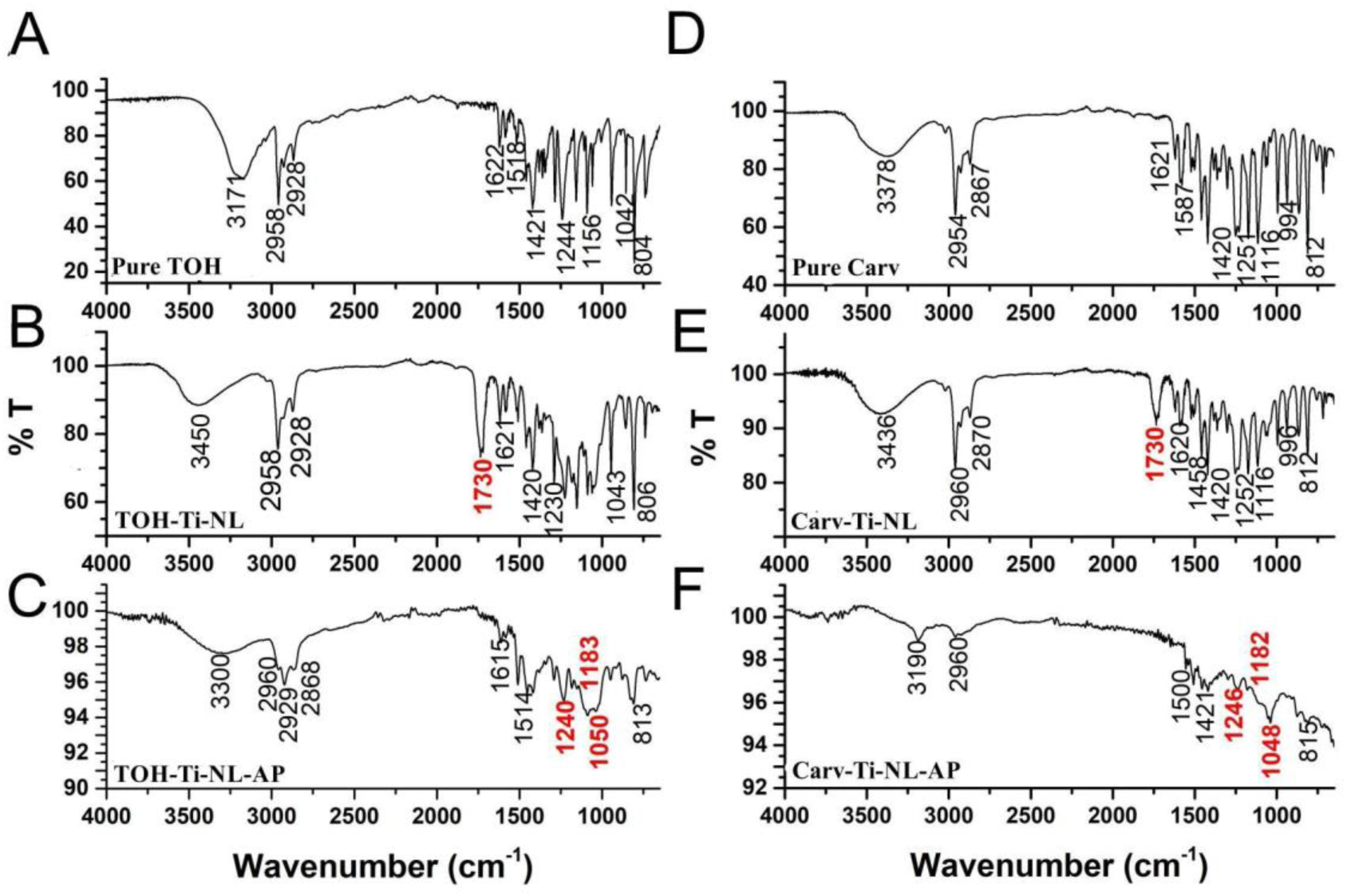
ATR-FTIR spectra of **(A)** pure TOH, **(B)** TOH-Ti-NL, **(C)** TOH-Ti-NL-AP, **(D)** pure Carv, **(E)** Carv-Ti-NL and **(F)** Carv-Ti-NL-AP.

After NLs formation, both compounds presented a similar broad peak at 3500-3400 cm^-1^ assigned to -HO participating in hydrogen bond between molecules and/or with the TiO_2_ surface. At the same time TOH-Ti-NL and Carv-Ti-NL presented a peak at 1730 cm^-1^, absent in the pure compounds, that indicates the spontaneous formation of ketonic groups that are also involved in the NLs formation on Ti surface. This peak is assigned to the C=O stretch due to the surface adsorbed species, where the oxygen adatoms are coordinated to Ti atoms [39]. The rest of peaks are present in the NLs, confirming the presence of aromatic ring and isopropyl groups.

##### 3.3.1.2 ATR-FTIR spectra of the NLs after anodic polarization

In order to investigate the chemical changes of the coatings by the electrooxidation process impelled by the AP in the −1.0V - +2.0V potential range, the FTIR analysis of the resulting NLs (TOH-Ti-NL-AP and Carv-Ti-NL-AP) was made, and the results are shown in Fig. 8E, 8F. The results showed a lower intensity in the FTIR signal, more notorious to Carv-Ti-NL-AP, depicting a thinner coating than before the treatment. TOH-Ti-NL-AP and Carv-Ti-NL-AP lost the ketonic signal (1730 cm^-1^ peak). A particular response was observed in case of TOH-Ti-NL-AP (Fig. 8F) showing strong signals at 1240 cm^-1^ and at 1050 cm^-1^, indicative of ether bond formation by dehydration reaction. This oxidized NL showed the -HO stretching signal displaced to 3300 cm^-1^, and it was assigned to a stronger adsorption to the surface. Significant changes in Carv-Ti-NL-AP spectra were detected (Fig. 8E). The band assigned to -OH stretching (3550-3200 cm^-1^) almost disappeared, just keeping a weak signal at 3190 cm^-1^. The typical contributions from the aromatic ring due to the C-H stretching (3100-2850 cm^-1^) was very weak and C=C stretching signal (1600-1500 cm^-1^) was almost undetectable. This NL mainly showed symmetric ether bond formation with strong signals at 1046 cm^-1^ and 882 cm^-1^. Therefore, these results showed different responses of the Carv-Ti-NL and TOH-Ti-NL, with Carv-Ti-NL being more susceptible to AP treatment than TOH-Ti-NL.

##### 3.3.1.3 ATR-FTIR spectra of the NLs the immersion in aqueous solutions

The ATR-FTIR analysis of the NLs after the release process in PBS medium was also made (Fig. 9). The spectra obtained after different release periods (1 h, 24 h, 48 h) revealed that in case of Carv-Ti-NL-R the 1730 cm^-1^ contribution attributed to ketone groups is weakened, as well as the peaks in the 3100-2850 cm^-1^ region (Fig. 9B). The longer the release period, the stronger the effect (see red arrows in Fig. 9B). Besides, in case of TOH-Ti-NL-R, after longer immersion periods, the large peak at the 3600-3100 cm^-1^ region becomes broader, depicting a higher signal at c.a 3300. All these effects suggest that the film degrades with increased immersion time, resulting in a noticeable reduction in transmittance. The ketonic group, although weaker, is detectable even after 48 h of immersion, indicating that these surface-bonded molecules are released from the NL, likely including unoxidized TOH and Carv. Also, the release of internal water from phenolic coatings is possible [40–42].

**Figure 9.**
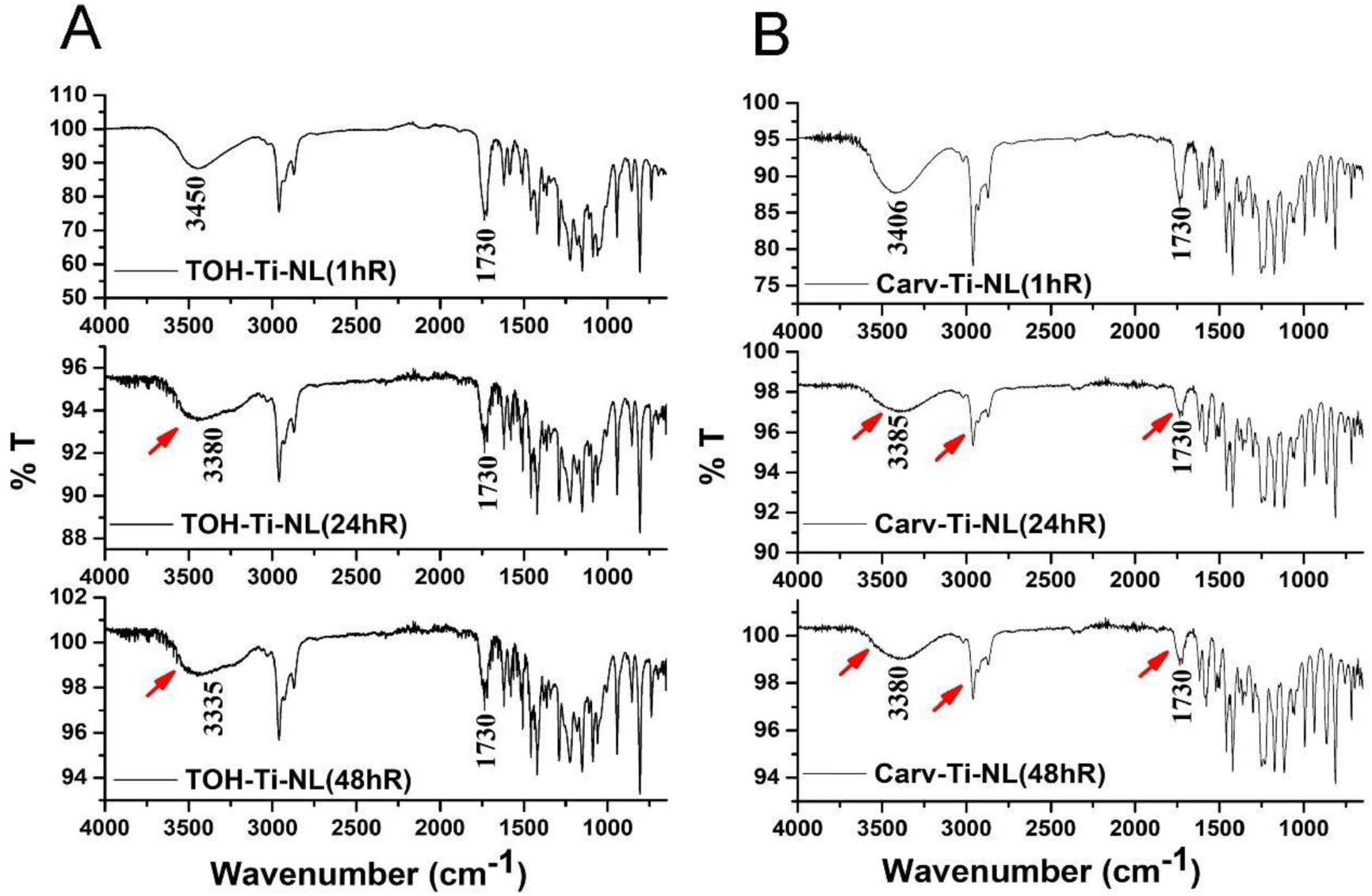
ATR-FTIR spectra obtained after different release periods in PBS solutions: 1, 24 and 48 h with TOH-Ti-NL (A) and Carv-Ti-NL (B) samples.

#### 3.3.2 Surface characterization by AFM: Roughness

The 2D AFM images (Fig. 10A-C above), show that both TOH and Carv form coatings on Ti with a close-packed structure. In Fig. 10D, the topographic profiles for the TOH-Ti-NL and Carv-Ti-NL samples are presented and analyzed in different zones: 1) before and after the step, which marks the boundary between covered and bottom areas, and 2) within the area covered by the NL. A noticeable difference emerges as the TOH-Ti-NL samples exhibit a profile with greater thickness variability in both the step and the film, while the Carv-Ti-NL samples demonstrate lower variability in both regions. The 3D images further emphasize this distinction, revealing that Carv-Ti-NL samples exhibit a more homogeneous and smoother NL.

**Figure 10.**
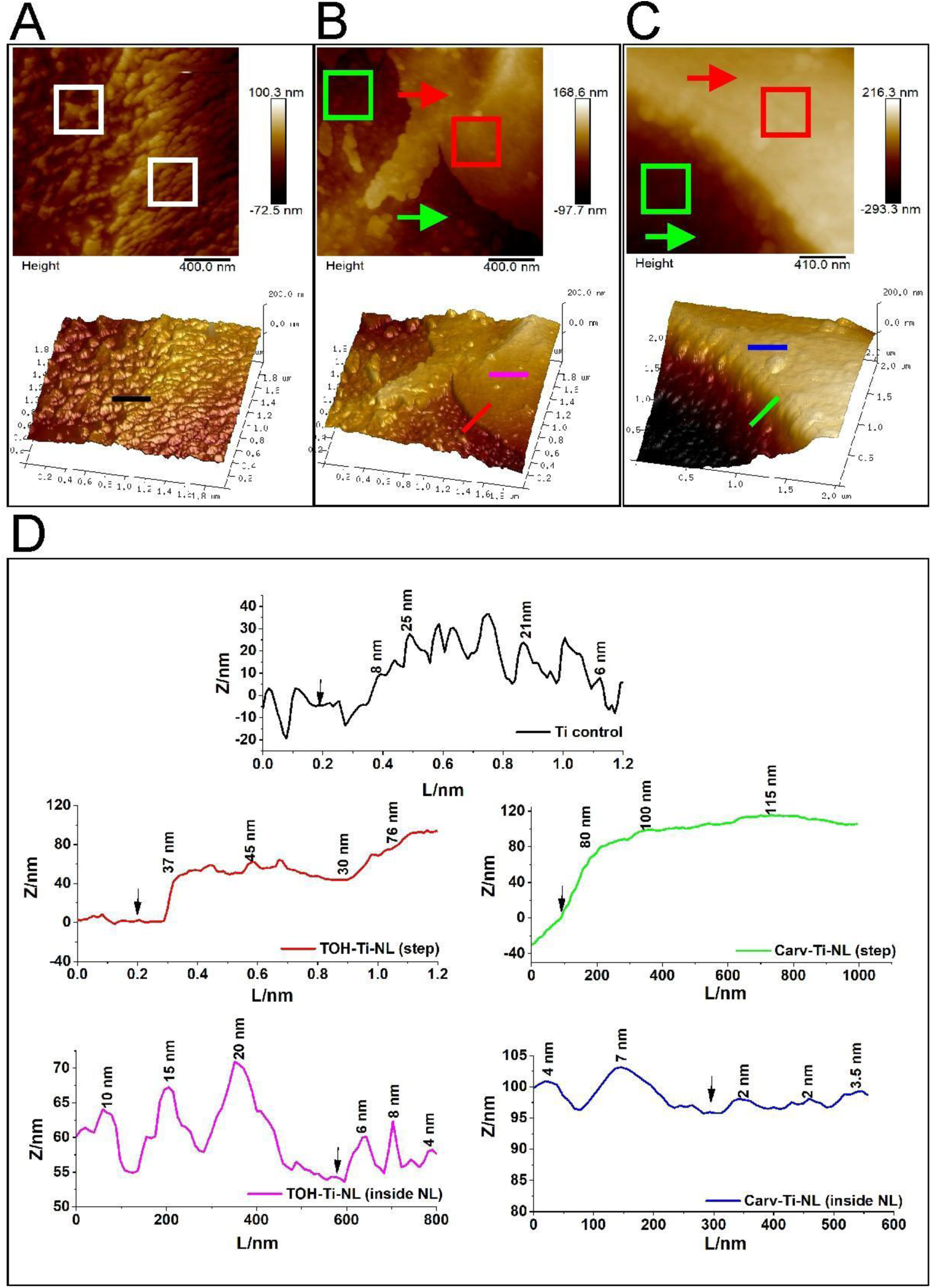
**2D** (above) **and 3D** (below) **topographic images obtained by AFM measurement** of **(A)** Ti control, **(B)** TOH-Ti-NL and **(C)** Carv-Ti-NL. **(D) Sections profiles** along the bars drawn in 3D images: Black for the control, red and fuchsia for TOH-Ti-NL and green and blue for Carv-Ti-NL. Black arrows indicate the reference zone for values calculated in the section’s profiles. Red arrows show areas covered by TOH or Carv NLs and green arrows show bottom areas. White, red and green squares indicate the zones where roughness parameters (Rq, Ra, Rmax, Rsa%) were obtained.

Red and fuchsia curves in Fig. 10D show the section profiles that are indicated with the colored bars in 3D AFM images (Fig. 10A-C below). It can be noticed that the Carv-Ti-NL coating is smoother (with distances between top and valleys of 1-5 nm) than that corresponding to TOH-Ti-NL. The roughness parameters of treated and untreated surfaces are summarized in Table 2. The parameters used to describe the surface were: root mean square roughness (Rq), average roughness (Ra), maximum roughness depth (Rmax), surface area relationship (Rsa), skewness (Ssk) and kurtosis (Sku). Rq, Ra and Rmax values, determined on the bottom areas of TOH-Ti-NL and Carv-Ti-NL samples, were slightly lower than those of the Ti control, although without any statistically significant differences (Table 2). However, when these parameters were obtained from the areas covered by the NLs, they were significantly lower than those of Ti control (p<0.001), indicating that the NLs flattened the topography of the surfaces. Accordingly, in presence of the NLs, Rsa values (2.1% for TOH-Ti-NL and 1.2% for Carv-Ti-NL) decreased significantly with reference to Ti control (16%). These results confirmed that the NLs smooth the topography of Ti surface with higher effect in case of Carv-Ti-NL (Rsa in whole image: 16%, 9±1% and 5±1% for Ti Control, TOH-Ti-NL and Carv-Ti-NL samples respectively).

**Table 2.**
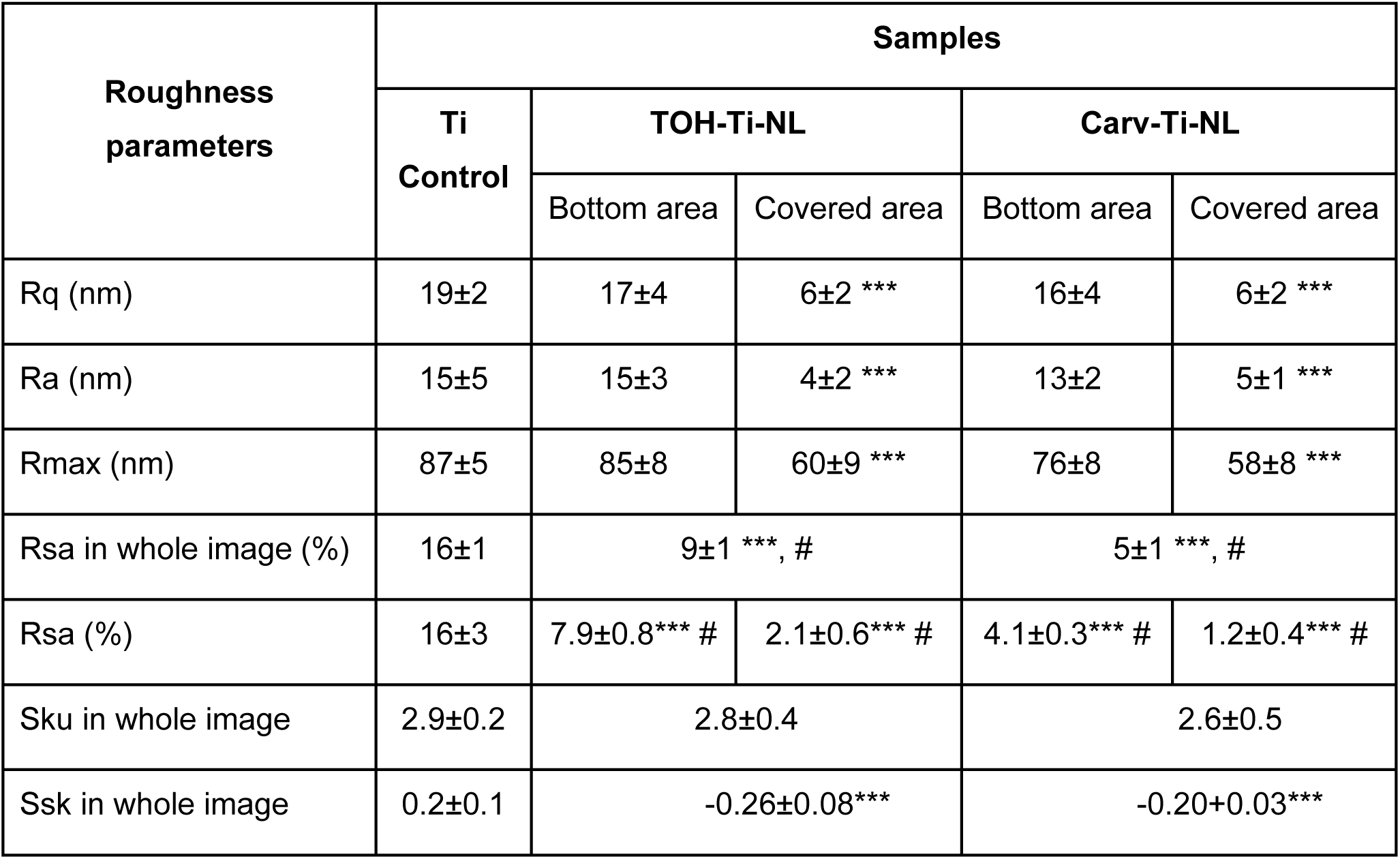
Roughness parameters from AFM image analysis of Ti Control, TOH-Ti-NL and Carv-Ti-NL images. *** Indicates significant differences respect to Ti control and # indicates significant differences between TOH-Ti-NL and Carv-Ti-NL (p<0.001).

| Roughness parameters | Samples |  |  |  |  |
| --- | --- | --- | --- | --- | --- |
|  | Ti Control | TOH-Ti-NL |  | Carv-Ti-NL |  |
|  |  | Bottom area | Covered area | Bottom area | Covered area |
| Rq (nm) | 19±2 | 17±4 | 6±2 *** | 16±4 | 6±2 *** |
| Ra (nm) | 15±5 | 15±3 | 4±2 *** | 13±2 | 5±1 *** |
| Rmax (nm) | 87±5 | 85±8 | 60±9 *** | 76±8 | 58±8 *** |
| Rsa in whole image (%) | 16±1 | 9±1 *** , # |  | 5±1 *** , # |  |
| Rsa (%) | 16±3 | 7.9±0.8*** # | 2.1±0.6*** # | 4.1±0.3*** # | 1.2±0.4*** # |
| Sku in whole image | 2.9±0.2 | 2.8±0.4 |  | 2.6±0.5 |  |
| Ssk in whole image | 0.2±0.1 | -0.26±0.08*** |  | -0.20±0.03*** |  |

Table 2 also shows Ssk and Sku values, important parameters to characterize implant surfaces [43]. Similar values were obtained for the three samples in case of Sku, but the sign changed in case of Ssk, positive for the control and negative for the covered Ti. Sku and Ssk values have not shown statistically significant differences between TOH- and Carv-Ti-NLs. It is noteworthy to mention that positive Ssk values indicate surfaces with sharper peaks and more rounded valleys, as in case of Ti control, whereas negative Ssk values suggest that peaks are more rounded, and valleys are sharper. Importantly, negative Ssk is associated with reduced sliding friction [44].

#### 3.3.3 Surface characterization: Wettability

The water contact angle (WCA) values measured on Ti control, TOH-Ti-NL and Carv-Ti-NL are shown in Fig. 2S. All the analyzed surfaces presented hydrophilic contact angles. Among them, TOH-Ti-NL exhibited a slightly lower hydrophilic behavior.

#### 3.3.4 Electrochemical behavior

##### 3.3.4.1 Open Circuit Potential measurements

The OCP vs time records for Ti samples with and without NLs immersed in solutions are shown in Fig. 11A. The OCP curve of the control depicts a continuous increase indicating that the passivation of the surface occurs due to the formation of additional TiO_2_ during the immersion time. Conversely, the curves of the Ti samples immersed in the 0.1M TOH or Carv in ethanol/sulfuric acid (30:70) solutions initially showed a sharp increase associated to the phenolic adsorption that leads to the formation of the NL. This increase is followed by a marked decrease thereafter to attain a nearly constant OCP value after c.a. 1000 s. However, at longer immersion periods, some instabilities in the OCP values were observed in case of TOH-containing solution (data not shown).

**Figure 11.**
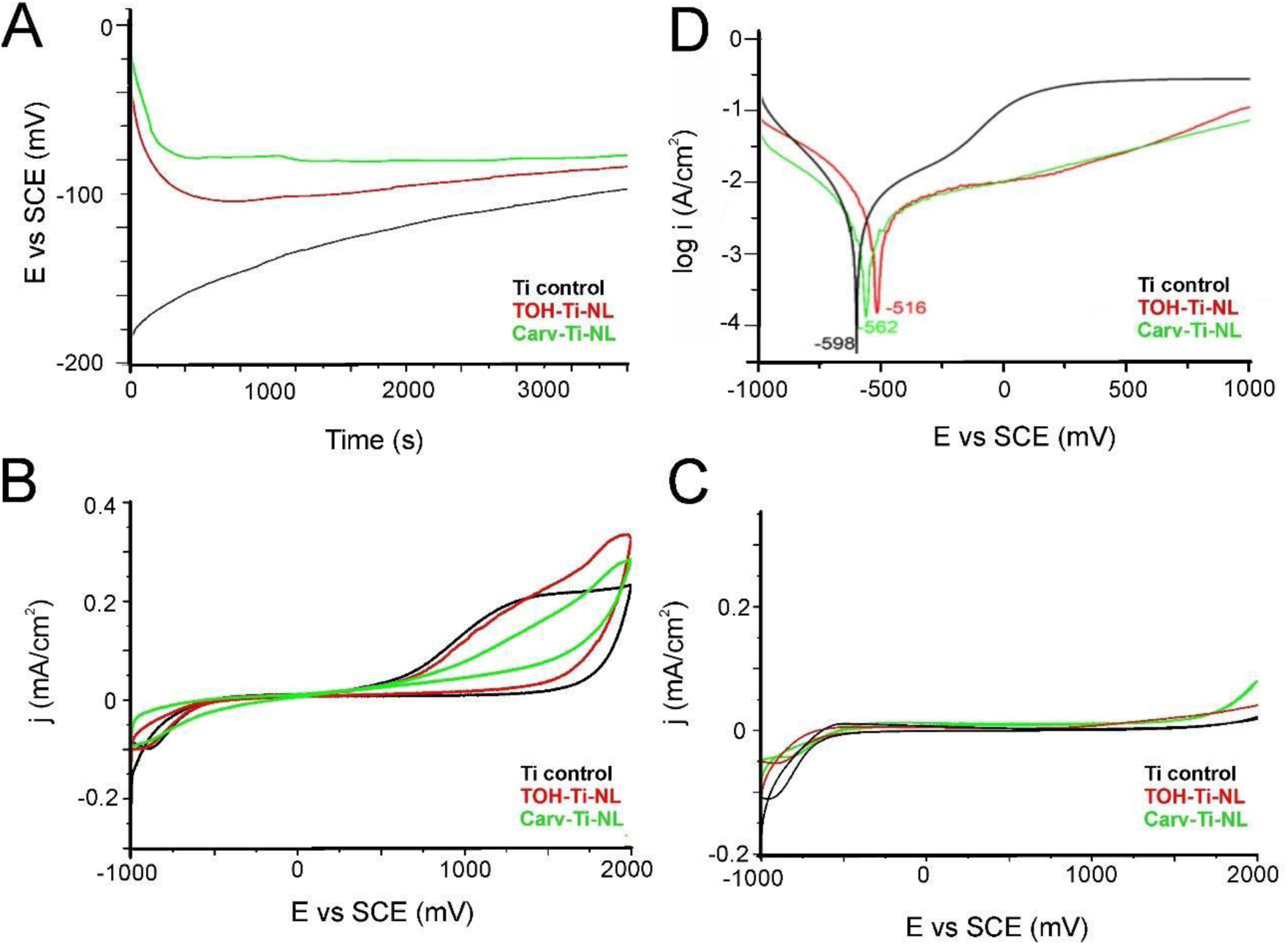
Electrochemical tests. **(A)** OCP measurements during TOH-Ti-NL and Carv-Ti-NL formation on Ti surface. Electrolyte: 0.1M TOH or Carv in ethanol/sulfuric acid (30:70) solution. For Ti control only ethanol/sulfuric acid solution was employed. **(B)** First cycle and **(C)** second cycle of cyclic voltammograms obtained for Ti control, TOH-Ti-NL and Carv-Ti-NL in 5 mM KCl solution at 50 mV/s within (−1000 mV, + 2000 mV) potential range. **(D)** Tafel curves obtained in 5 mM KCl at 1 mV/s.

##### 3.3.4.2 Cyclic voltammetry

The electrochemical behavior of TOH- and Carv-Ti-NL was also investigated by cyclic voltammetry (CV) in the (−1 V, at +2 V) potential range at 50 mV/s using 5mM KCl as electrolyte. Fig. 11B shows the voltammogram of the first cycle of Ti control, TOH-Ti-NL and Carv-Ti-NL voltammogram while Fig. 11C shows the second cycle. The voltammogram of Fig. 11B shows that at potentials higher than 0.5 V, an increase in the current density can be noticed in the Ti control voltammogram due to the formation of additional oxide on Ti. In case of the curves of TOH-Ti-NL, during the anodic scan of the cycle, higher currents densities than the control were recorded due to simultaneous formation of TiO_2_ and the electro-oxidation of TOH-Ti-NL coating. In case of Carv, less TiO_2_ seems to be formed on the surface and Carv-Ti-NL oxidation occurs. The reverse scan showed that the cathodic currents were lower than the anodic ones, being higher in case of Carv layer due to the additional electrooxidation of the Carv-Ti-NL layer. In view that the anodic contributions related to the oxide formation are not accompanied by the corresponding cathodic reaction it can be assumed that irreversible redox processes occur.

##### 3.3.4.3 Tafel curves

Tafel curves were obtained using a KCl solution (5 mM) as electrolyte. Fig. 11D shows that TOH-Ti-NL and Carv-Ti-NL samples depict markedly lower anodic current values than the polished Ti sample at potentials more positive than Ecorr. They also show more anodic Ecorr values (Table 3) due to the presence of the adsorbed layers.

**Table 3.** Values of Ecorr and Icorr obtained from Tafel curves in 5 mM KCl.

| Samples | E<sub>corr</sub> (mV) | I<sub>corr</sub> (μA/cm<sup>2</sup>) |
| --- | --- | --- |
| Ti control | -598 ± 4 | 38 ± 5 |
| TOH-Ti-NL | -516 ± 3 | 210 ± 2 |
| Carv-Ti-NL | -568 ± 5 | 150 ± 2 |

## 4. DISCUSSION

In this study, two NLs based on phenolic monoterpenes isomers (TOH and Carv) were prepared using a simple one-step immersion treatment method on Ti samples. The biological response of bacteria and eukaryotic cells was assessed, and a comparative analysis of the physicochemical and surface properties of the coatings was conducted. The aim of this analysis was to elucidate potential physicochemical differences (evaluated by ATR-FTIR, electrochemical analysis, AFM roughness assessment and wettability measurements) that could distinctly impact the antibacterial behavior and potential osseoinductive effect of the surfaces. The main factors involved in these behaviors, along with information previously reported by other authors, will be discussed in the following sections.

### 4.1 Similarities and differences in NLs formation and properties

#### 4.1.1 NLs’ formation mechanism

The ability of TOH and Carv to form hydrogen bonds through the -OH group on the Ti surface and/or other molecules, along with the Van der Waals forces, are mainly responsible for the adsorption process on Ti, as previously observed with several substrates [45–47]. The formation of self-assembled layers is also supported by QSAR analysis, which showed that Carv and TOH exhibit low free energy of solvation and dipole moment. This may lead to a higher likelihood of accumulation of these organic molecules on metal surfaces. Moreover, both compounds share similar adsorption energy and interaction distance with the surface [48,49].

According to the FTIR results (Fig. 8), both NLs showed similar spectra. Hence, it can be hypothesized that the coating mechanism in both NLs was similar and is as follows: Firstly, molecules of TOH and Carv are adsorbed through hydrogen bonding of -OH groups onto the TiO_2_ surface. Subsequently, spontaneous oxidation reactions occurred after the adsorption process. A new peak at 1730 cm^-1^ revealed the formation of a keto group in both NLs. It is assigned to a coordination between the ketone group and the Ti^+4^ ion from the TiO_2_ surface [39]. In agreement, keto groups were also detected on other metals such as Cu surfaces after the spontaneous adsorption of TOH and Carv [47]. However, it can be inferred that some molecules are not affected by this oxidation process, as the -OH signal is also detected in the spectra. These unaltered molecules of TOH and Carv remain within the NLs, interacting mainly through hydrogen bonding with other molecules and TiO_2_ surface [47,50].

In line with our observations, Mangiacapre *et al* [51] have recently investigated Carv and TOH in liquid state (at high temperature). The authors demonstrated that for Carv, there is a tendency for the aromatic rings to orient perpendicularly. In contrast, TOH molecules exhibit a more heterogeneous structural organization characterized by both parallel and antiparallel orientations of neighboring rings. TOH preferentially forms hydrogen bonds between its hydroxyl groups. Conversely, Carv demonstrates a stronger inclination towards O-H···ㄫ hydrogen bonding interactions. It was considered that Carv arranged in a more structurally organized manner than TOH. Therefore, the distinct tendency of Carv and TOH to molecular arrangements have the potential to influence the physicochemical characteristics of NLs.

#### 4.1.2 Roughness and wettability

Differences in roughness and thickness of the NLs in this work were observed through 3D images and section profiles (Fig. 10) from AFM images. Carv-Ti-NL showed a more uniform, smoother, and compact coating than TOH-Ti-NL. The analysis of roughness parameters suggests that, although Ra, Rq and Rmax are commonly used to characterize surfaces, being able to detect differences between the control and the samples with NLs, they lack sensitivity to detect variability between Carv-Ti-NL and TOH-Ti-NL. Conversely, Rsa (%) emerged as a relevant parameter in the roughness analysis of these coatings that fit with the AFM images. Results reveal that Rsa values were smaller and, consequently, samples with NL are smoother than that of Ti control with the smallest value for the smoothest sample, Carv-Ti-NL. All these data revealed that the surface topography is flattened by the organic NLs and the Carv-Ti-NL was significantly smoother than TOH-Ti-NL.

The sign of Ssk was positive in case of control Ti and negative for the covered samples. It describes the Ti control surface having sharper peaks and rounded valleys, while more rounded peaks with sharper valleys characterize the surfaces with NLs. The negative values are associated with more sliding surfaces and sometimes encourage cell attachments and colonization.

The differences observed in roughness did not imply significant differences in wettability (Fig. 2S). The measured WCA indicated that all surfaces studied were hydrophilic. However, only the TOH coating showed a slightly increased value of WCA compared to the Ti control. Carv samples exhibited similar WCA values to the Ti control. Similar results were found in self-assembly of chitosan+caffeic acid on Ti alloy, where increasing roughness resulted in moderate hydrophilicity [52].

#### 4.1.3 Electrochemical behavior

It is known that the TiO_2_ layer formed on the metal surface exhibits a duplex structure, which is comprised of a dense inner layer and a porous outer layer [53]. According to previous reports, results showed a continuous and slow increase of the OCP during the first hour for Ti control after the immersion in solutions without PhCs attributed to the formation of additional dense inner oxide layer [33,54]. In case of the immersion of the Ti samples in the PhC solutions, the development of the adsorbed organic layer is denoted by the abrupt increase of the OCP [54–57]. Tafel curves recorded during the anodic scanning showed lower currents at potentials more anodic than Ecorr, indicating that lower amounts of oxide are formed on Ti surface when the NLs are present.

During CV assays, the first cycle revealed the oxidation of the PhCs in the oxide region (+0.5V - +2.0V) during both the anodic and cathodic scans. A dissimilar response of Carv-Ti-NL and TOH-Ti-NL in aqueous solutions was revealed by CV (Fig 11B-C) since different oxidation currents were detected for the two NLs. However, it was reported that when other electrolyte such as acetonitrile is used, the oxidation of TOH and Carv depict analogous anodic contributions indicating that the electrochemical response is dependent on the solvent and revealing the dissimilar interaction of Carv-Ti-NL and TOH-Ti-NL with water [58]. The second cycle of CV assays (Fig. 11C) showed that the remaining oxidized NLs blocked the metal surface and impeded additional oxide formation, since very low current densities in the +0.5 V – +2.0 V potential range were recorded. Similar behavior was reported for other metal substrates [46,47,59]. Interestingly, after the anodic polarization (AP) of TOH-Ti-NL and Carv-Ti-NL, FTIR spectra showed marked dissimilitude. Particularly, the 3300 cm^-1^ contribution is only present in case of TOH-Ti-NL-AP. Consequently, it can be inferred that TOH-Ti-NL and Carv-Ti-NL, although showing a similar FTIR spectra before the AP, have different chemical properties that lead to distinct oxidation products detected by FTIR analysis after the AP.

#### 4.1.4 Release profiles

According to Fig. 1, which depicts the cumulative concentration of the released compounds in the solution, TOH-Ti-NL demonstrates a distinct profile with a higher release compared to Carv-Ti-NL throughout the entire assay period.

It is noteworthy that within the initial 12 h, there is a high release rate (frequently known as burst release) of both compounds. This can be attributed to the release of the previously mentioned molecules of TOH or Carv trapped within the NLs, which are immediately liberated upon contact with the aqueous solution. However, the presence of undetectable quantities of their oxidized keto derivatives should not be disregarded [60], as the main groups characterizing these molecules were found in the NLs (Fig.8B,8E). The concentration *vs* time curve over the first 24 h-release period can be simulated with a kinetic equation derived from the modified Korsmeyer-Peppas model [61]. The application of this model to our data allows us to conclude that the Carv release exhibits a slower rate with a longer lag time than that of TOH. These findings suggest that the distinct physicochemical properties of both NLs lead to different release profiles.

After 24 h, Carv was released at a constant rate up to 10 days, whereas TOH-Ti-NL released higher amounts of the compound in three well-defined steps. In line with this data, it is important to consider that the solubility in water (980 µg/mL for TOH and 830 µg/mL for Carv) is higher for TOH and contribute to the driving force behind the spreading of the compound from the film towards the water medium [62]. Moreover, as mentioned before, other physicochemical properties of the NLs may also influence the observed release profiles. Carv release profile appears to be continuous and associated to the homogeneous structure of the NL (Fig 10, table 2) whereas the release behavior of TOH-Ti-NL (in three steps) may be related to its irregular surface structure (Fig 10B) [63]. Thus, after 24 h immersion, “patches” of TOH-Ti-NL may absorb water intermittently, causing a cyclic pulse of patch hydration and compound release, which is consistent with the three-step profile shown in Fig. 1. Conversely, the release profile of Carv-Ti-NL can be attributed to the gradual hydration of the compact layer, leading to a continuous release of the compound [51,64]. Supporting these hypotheses, the ATR-FTIR analysis (Fig 9) of the NLs after the release process, revealed that the spectra were similar to those corresponding to the recently prepared NLs but with weakened signals.

Furthermore, considering Fick’s second law equation to describe and simulate the variation of compound concentration with location and time from a source [65] it can be inferred that the concentration of TOH and Carv at the NL surface is several times higher than the average concentration measured and reported in the release profile. This high surface concentration crucially impacts the biological response.

### 4.2 Impact of the physicochemical parameters of NLs on the biological systems. A comparative analysis

#### 4.2.1 Antimicrobial effect of NLs

It was reported that higher bacterial cell attachment was found on surfaces with negative Ssk values than on those of positive Ssk and flat surfaces [44]. However, our results do not show this tendency indicating that other physicochemical properties of the NLs, particularly the release parameters, appear to impact bacterial adhesion and viability more than the topological aspects.

Carv and TOH, can function as proton exchangers, thereby reducing the pH gradient across the cytoplasmic membrane, leading to the collapse of the proton motive force and depletion of the ATP pool [66]. The positioning of the -OH group of the isomers seems not to be decisive for the antimicrobial efficiency [14]. Importantly, the toxicity of TOH and Carv is strongly linked to the alteration of *quorum* sensing (QS). In presence of phenolic compounds, a significant reduction in the production of QS autoinducers and inhibition of motility was detected [7,67].

Considering the diameter of the inhibition halos (Table 1) it becomes evident that the compounds from Carv-Ti-NL and TOH-Ti-NL diffuse through the agar surface, impeding bacterial growth in the vicinity of the samples. Based on the Carv and TOH release measurements (Fig 1), it can be concluded that the compounds are released by the NLs both in aqueous media and in a gelled medium (Table 1). Moreover, no significant differences in antimicrobial effect between both compounds are observed within the first 24 h (Table1, Fig. 2, Fig. 3).

However, as previously mentioned, it is important to consider that the concentration of active principles at the coating/bacteria interface may be significantly higher than the concentration in the bulk solution measured during the release assays [65]. Therefore, due to concentration gradient and burst release effect there is a stronger impact on the cells attached at the interface compared to those farther away. Results showed that the average concentrations in PBS after 24 h of release were approximately 20 and 17 µg/mL for TOH and Carv, respectively. Although apparently small, these values are associated with higher concentrations at the NL surface that exhibit great efficacy as antimicrobial agent during the first 24 h, effectively killing the bacteria attached to the surface, preventing the evolution of biofilms and leading to an eradication effect [68]. Consistently, present results depict the reduction of EPM from *S. aureus* induced by both NLs (Fig. 4) in line with prior studies [67]. This decrease implies that the formation of colonies is hindered leading to isolated bacteria that are more labile to the antimicrobial agent attack [26,69–72]. In agreement with our results, Ti surfaces modified with Carv and TOH-containing essential oil, incorporated into plasma-sprayed hydroxyapatite coating, provide infection prevention against *S. epidermidis* in load-bearing orthopedic and dental applications [73].

After the initial 24 h-burst release period, Carv is released linearly at a lower rate than TOH, which exhibits a sharp increase. Despite this difference in the release profiles, a lower but still very significant bactericidal effect (4 orders of magnitude decrease in CFUs) is observed in both cases after 24 h of release (NL-(24hR) (Fig. 2, Fig. 3). After 48 h of release (NL-(48hR)), the NLs lost their antimicrobial activity because the surfaces become depleted after 48 h, likely falling below the MIC value. Under these conditions, bacteria can attach, proliferate and produce significant amounts of EPM, resulting in biofilm formation (Fig. 4). This is consistent with previous reports of significant bacterial attachment on metal areas with very low antimicrobial concentration [46].

#### 4.2.2 Effect of the NLs on the attached pre-osteoblastic and fibroblastic cells

TOH-Ti-NL exhibits distinct behavior compared to Carv-Ti-NL when interacting with eukaryotic cells. Specifically, TOH-Ti-NL demonstrated a significant increase in osteogenic cell proliferation without concurrent enhanced fibroblast proliferation (Fig. 5). On the contrary, the results revealed that Carv-Ti-NL samples displayed a higher affinity for fibroblastic cells than for pre-osteoblastic cells. The analysis of the parameters that could influence this response is subsequently made.

##### 4.2.2.1 Topographic surface properties

Contrary to the impact of NLs on bacteria, differences in topographic characteristics seem to directly affect the attachment of eukaryotic cells in a manner contingent upon the specific cell line. As we mentioned previously, significant differences in roughness were observed between the two NLs. Carv-Ti-NL presented a smoother and more compact appearance, as revealed by Rsa% parameter. In this context, several authors have presented findings on various surfaces that align with our own. Benz *et al*. [74] highlighted a factor contributing to the robust adhesion of osteoblasts to Ti, attributing it to the presence of –OH groups on the surface that facilitate the development of an apatite structure, thereby promoting the adhesion and growth of osteoblasts. Contrastingly, a coating with polyetheretherketone (PEEK), known for its exceedingly smooth, chemically inert, and hydrophobic surface, exhibited poor osteoblast adhesion. Conversely, a relatively irregular surface resulting from a coating with Poly-L-Lysine (PLL) encouraged osteoblast adhesion. Similarly, titanium surfaces with anisotropically patterned dense nanospikes have been shown to promote osteoblast activation [75]. It is noteworthy that extremely smooth or excessively rough surfaces do not seem to offer advantages for osteoblast growth, underscoring the sensitivity of this cell line to surface roughness. Conversely, fibroblasts on the smooth PEEK surfaces demonstrated widespread distribution, with no adverse effects on their cellular growth. Previous research has indicated a high proliferation rate for epithelial cells and fibroblasts on smooth surfaces [76] or surfaces with nano-roughness [77] where sliding processes are favored. Kunzler *et al* [78] speculate that osteoblast proliferation is preferred on the locally smooth but macroscopically rough surface structures. Experiments with rat calvarial osteoblasts (RCO) and human gingival fibroblasts (HGF) on a gradient of roughness on aluminum also demonstrated that RCOs showed a significantly increased proliferation rate with increasing surface roughness while fibroblasts showed decreased proliferation on rougher substrata [78]. Some types of roughened surfaces have also been found to produce better *in vivo* osseointegration than smoother surfaces, suggesting that the surface modulates the bone response including osteoblast differentiation, extracellular matrix deposition, and calcification.

Our results are consistent with all these previous studies. Thus, cells grown on TOH-Ti-NL (more irregular, rougher and heterogeneous than Carv-Ti-NL) showed significantly higher osteogenic markers than on Ti control (the roughest surface in this work) or Carv-Ti-NL (the most compact and smoothest coating), revealing the sensitivity of this cell line to surface roughness (Fig. 7, Fig. 10) [79,80].

##### 4.2.2.2 Cytotoxicity of the compounds

Despite the primary influence of topological properties on the adhesion and proliferation of various cell lines, other factors such as the cytotoxicity of the compounds need to be taken into consideration.

The MTT assay with 24 h-extracts from NLs on fibroblasts and pre-osteoblasts (Fig. 6) were made. From the release curve (Fig. 1), the concentrations of TOH and Carv in the extracts (after 24 h of release) is estimated as c.a. 20 and 17 µg/mL, respectively. Results showed that neither TOH-Ti-NL nor Carv-Ti-NL extracts caused any cytotoxic effect on fibroblasts, while pre-osteoblasts exhibited a significant decrease in cell viability in the presence of the Carv-Ti-NL extract. Consistent with our results, lower cytotoxicity of TOH compared to Carv has been reported in other cell lines. In CHO-K1 cells, usually used as a model to evaluate cytotoxic effects of drugs, the concentration at which viability is less than 70% was reported as 75 and 25 µg/mL for TOH and Carv, respectively [81]. Thus, TOH appears to have lower cytotoxicity in CHO-K1, L929 fibroblasts, and MC3T3 pre-osteoblasts, while Carv has been shown to be cytotoxic in CHO-K1 and MC3T3. L929 fibroblasts seem to be resistant to Carv.

Other authors have presented findings on various surfaces and their analysis and interpretations align with our own [44,82,83]. These observations indicate that the biological effects of these compounds depend, not only on the morphological characteristics of the coatings they are part of, but also on their intrinsic toxicity.

Overall, our results showed that TOH-Ti-NL promoted a significant increase in osteogenic cell proliferation *in vitro* without concurrent enhanced fibroblast proliferation. Studies with thymol-loaded on chitosan hydrogel exhibits both, *in vitro* and *in vivo,* improved osteogenic activity [84] This characteristic makes it a promising candidate for application in orthopedic or dental implants with an additional highly effective antimicrobial action. In contrast, fibroblast displayed higher affinity for Carv-Ti-NL and may induce an undesirable fibrous encapsulation process at the implant surface avoiding the osseointegration [85]. These results suggest that Carv-Ti-NL may find utility in other applications where the biomaterial needs to be in contact with the gums, making them potentially valuable for dental abutments. Considering the individual properties of each NL, one might select the use of either Carv-Ti-NL or TOH-Ti-NL in order to adjust specific chemical physical properties of interest for biological systems.

### Conclusions

Carv-Ti-NL and TOH-Ti-NL were formed on Ti surface using a straightforward one-step immersion treatment technique. The chemical and physical properties of Carv-Ti-NL and TOH-Ti-NL were evaluated to elucidate their relationships with biological effects. The ATR-FTIR spectra of the NLs were similar and indicated the spontaneous oxidation of Carv and TOH leading to ketonic structures; however, distinct contributions were observed after the electrooxidation of each NL, linked to the different molecular interactions of organic molecules with each other and with the surface. Slight differences in hydrophilicity were found for both NLs, but significantly higher value of Rsa roughness parameter was found for TOH-Ti-NL. Furthermore, the release curves of Carv and TOH from the NLs revealed distinct profiles over time, with higher release in case of TOH-Ti-NL.

Despite differing physical and chemical characteristics, both TOH-Ti-NL and Carv-Ti-NL showed high antimicrobial efficacy on Ti surfaces, eradicating bacteria within 24 h. This efficacy is attributed to the high surface concentration of TOH and Carv at initial times and the burst release effect. The bactericidal effect persisted until 48 h, making them promising to prevent infections associated with Ti implants. However, after a 48-h release period, antimicrobial activity ceased due to TOH and Carv depletion. Nevertheless, their effectiveness within 48 h post-implantation is significant, ensuring microbial eradication beyond surgery.

The roughness of NLs appears to distinctively influence the attachment and metabolism of the assayed eukaryotic cell lines. Results demonstrate that under *in vitro* conditions, pre-osteoblasts exhibited a preference for TOH-Ti-NL (rougher than Carv-Ti-NL) whereas fibroblasts thrived better on smoother NLs, as in the case of Carv-Ti-NL, characterized by the lowest Rsa% value. TOH-Ti-NL did not have a cytotoxic effect on pre-osteoblast and fibroblast cells, while Carv-Ti-NL extract showed cytotoxicity on pre-osteoblastic cells.

The comparative analysis of the NLs formed with Carv and TOH isomers leads to conclude that the enhanced osteogenic activity and the strong antimicrobial effectiveness of TOH-Ti-NL render it promising for bone-related medical applications, while Carv-Ti-NL are suitable for promoting fibroblast growth. Both coatings offer tailored chemical and physical properties for diverse biological systems, suggesting versatile applications.

## Acknowledgments

This work was supported by CONICET [PIP 2021 11220200100315CO, doctoral fellowship of AG]; ANPCyT [PICT 2019-00631, PICT-2020-02169, PICT Start Up 2020-0034]; and UNLP [Project 11/X900]

## Declaration of Generative AI and AI-assisted technologies in the writing process’

During the preparation of this work the author(s) used ChatGPT in order to check grammar, spelling and to improve readability of the text. After using this tool/service, the author(s) reviewed and edited the content as needed and take(s) full responsibility for the content of the publication.

