## Supplementary information for "Antimicrobial nanolayers of thymol and carvacrol on titanium surfaces: the crucial role of interfacial properties in thymol’s superior osteogenic response"

1. **UV-Vis spectroscopy of TOH and Carv solutions**

The UV-Vis spectra of TOH and Carv solutions were measured using a Shimadzu UV-1800 spectrophotometer in a quartz cuvette. Quantification of TOH and Carv released from the nanolayers (NLs) were performed by comparison with a calibration curve.

UV-Vis spectra of Carv and TOH solutions are shown in Fig 1S. A single sharp band at 274 and 273 nm were found for TOH and Carv, respectively.

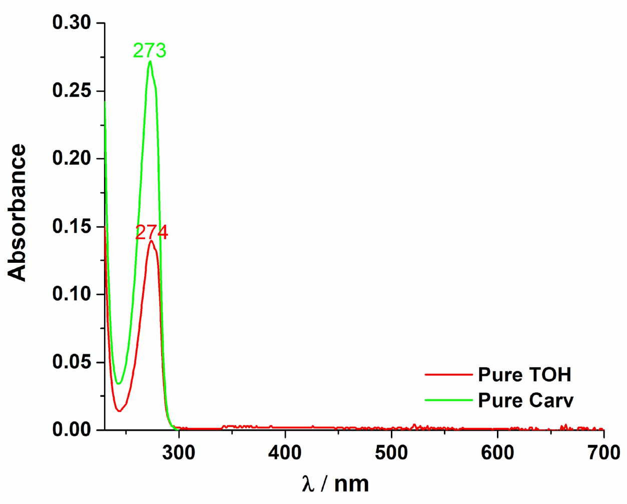


**Figure 1S.** UV spectrum of pure thymol and carvacrol

2. **Water contact angle measurements (WCA)**

The water-in-air contact angle (WCA) was measured on the Ti control, TOH-Ti-NL and Carv-Ti-NL. WCA measurements were obtained by dispensing 1 μL of distilled water onto each surface. All measurements were conducted at room temperature using an automated goniometer (Ramé-Hart Instrument co. Mod 290-U1) [1].

The results of WCA measurements (Fig. 2S) showed that all studied sufaces were hydrophilic. However, differential behavior was found between TOH-Ti-NL and Carv-Ti-NL. TOH-Ti-NL showed a slightly increased WCA value ($p<0.001$) compared to Ti control, while Carv samples exhibited a WCA value similar to that of the Ti control but lower than that of TOH-Ti-NL ($p<0.001$).

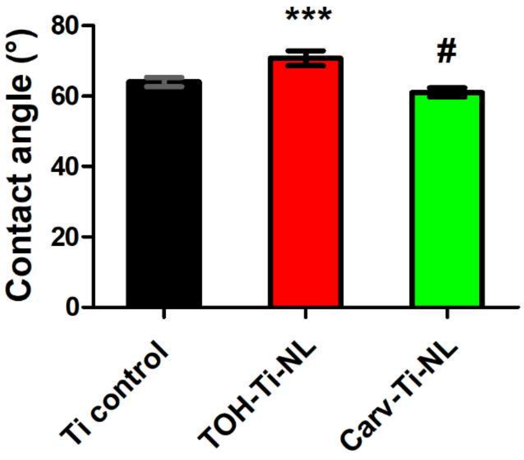


**Figure 2S. Water Contact Angle measurements**. *** Indicates statistically significant differences respect to Ti control and # indicates statistically significant differences between TOH-Ti-NL and Carv-Ti-NL ($p<0.001$).

**3. Burst release model**

The release process from the TOH-Ti-NL and Carv-Ti-NL during the first 24 h in PBS solution, can be simulated with a kinetic equation derived from the modified Korsmeyer-Peppas model [2]:

$$\mathrm{Mt}\ (\%) = \frac{a\ (t-tL)}{1+b\ (t-tL)}$$

where Mt% is the percentage of drug released at time t, $a$ (% $min^{-1}$) and $b$ ($min^{-1}$) are the characteristic parameters of the model and tL is the lag time, which is the value of time obtained by extrapolation of the model for Mt (%) = 0. The following data were obtained from the simulated curve applied to the experimental data obtained in this work: $a_{Carv}$= 1.7; $a_{TOH}$= 3.0; $b_{Carv}$= 0.056, $b_{TOH}$= 0.098; $t_{Lcarv}$= 1.0 h, $t_{LTOH}$ = 0.5 h.

This allows us to infer that the kinetics of Carv release show a slower rate (with lower *a* and *b* parameters) than TOH and a longer tL. However, the model only applies up to 24 h of release, during the burst release associated with the efficient bacteria eradication effect, which is similar for both Carv and TOH.
